# Targeting glucocorticoid signalling in dendritic cells for glioblastoma treatment

**DOI:** 10.64898/2026.08.07.742730

**Authors:** Qiuchen Zhao, Jhuma Pramanik, Arif Jabbar, Yifan Gui, Soura Chakraborty, Emily Fysh, Subhanita Ghosh, Pilar Puerto-Camacho, Miguel M. Santos, Aws Al-deka, Matthaios Pitoulias, Hosni Hussein, Christopher J. Ward, George Gentsch, Natalie Z. M. Homer, Benjamin Czech Nicholson, Rahul Roychoudhuri, Richard Mair, Bidesh Mahata

## Abstract

Glioblastoma (GBM) remains a devastating disease with few meaningful therapeutic advances over the past three decades. Dendritic cell (DC) vaccination is a promising immunotherapeutic strategy for GBM, but its efficacy is limited by the clinical use of dexamethasone to control cerebral oedema and associated symptoms. Here we show that steroid signalling is a central regulator of DC dysfunction in GBM. Through targeted metabolomics of primary GBM samples, we identified a steroid-rich tumour microenvironment in which dexamethasone is present at high levels. Across bulk and single-cell transcriptomic and epigenomic datasets, NR3C1 emerged as the dominant steroid receptor in GBM immune cells and was negatively associated with activated DC states. In patient-derived DCs, dexamethasone altered NR3C1 chromatin occupancy, induced broad transcriptional and chromatin remodelling, and suppressed co-stimulatory antigen-presentation and cytokine programmes. DC-specific deletion of Nr3c1 restricted syngeneic glioblastoma growth, enhanced DC activation, promoted cytotoxic CD8^+^ T cell responses and remodelled myeloid states in vivo. In GBM patient-derived DCs, pharmacological or non-viral CRISPR-mediated disruption of NR3C1 restored inflammatory, antigen presentation and T cell-stimulatory programmes, enhanced antigen-specific CD8^+^ T cell priming, and improved tumour lysate-loaded DC vaccination. Together, these findings identify glucocorticoid signalling as a key barrier to DC immunotherapy in GBM and establish NR3C1-targeted, steroid-resistant DCs as a potential therapeutic strategy.

## Introduction

Glioblastoma (GBM) remains the most aggressive primary brain tumour in adults and is associated with a dismal prognosis despite maximal safe resection followed by radiotherapy and temozolomide, a treatment backbone that has changed little since its establishment in 2005^1,2^. Although this treatment protocol improves survival, long-term outcomes remain poor and no curative therapy exists^1–4^. Efforts to translate immunotherapy into GBM have likewise been disappointing: phase III trials of PD-1 blockade and other vaccine-based approaches have failed to improve survival, underscoring the profound immunosuppressive and therapy-resistant nature of the GBM microenvironment^5–11^. Among immunotherapeutic strategies, dendritic cell (DC) vaccines remain particularly attractive because they directly engage antigen presentation and T cell priming, and several clinical studies have reported encouraging survival signals^12–15^. However, their efficacy has remained variable, and the mechanisms that limit DC function in patients with GBM remain incompletely defined.

Glioblastoma develops within a profoundly immunosuppressive tumour microenvironment, in which myeloid dominance, defective antigen presentation and dysfunctional lymphocyte states together limit effective anti-tumour immunity^16,17^. Steroid signalling is increasingly emerging as one such suppressive axis. Previous studies have shown that steroids can be synthesised, recycled or amplified locally within tumours and tumour-infiltrating immune cells, where they promote CD8^+^ T-cell dysfunction, regulatory T cells (Treg)-mediated suppression and dendritic-cell dysregulation^18–24^. Consistent with this concept, we and others recently identified local steroid and steroid signalling programmes in human solid tumours and showed that genetic or pharmacological perturbation of steroid pathways can restore anti-tumour immunity and restrain tumour growth^19,25,26^. Importantly, exogenous synthetic steroids, such as dexamethasone, remains very widely used in patients with GBM to control peritumoural cerebral oedema^27^, yet accumulating clinical and preclinical evidence links its use to poorer survival, a more immunosuppressive glioma microenvironment and reduced benefit from immunotherapy^28^. Although this connection has not been fully resolved for DC vaccination specifically, the potential relevance of dexamethasone to vaccine efficacy is supported by a recent personalised neoantigen vaccine trial in GBM, in which robust vaccine-induced T-cell responses were observed in patients who were not receiving dexamethasone during vaccination, whereas concurrent dexamethasone exposure was associated with markedly reduced immunogenicity^29^. Together with the association of lower concurrent dexamethasone exposure with better outcome in a recent GBM DC vaccine study^30^, these findings support that steroid signalling may represent a clinically relevant barrier to effective DC immunotherapy in GBM.

DC vaccination is an attractive strategy in GBM because it seeks to overcome defective endogenous priming by generating autologous monocyte-derived DCs (moDCs) *ex vivo*, loading them with tumour lysate or defined antigens, and reinfusing them to initiate tumour-specific T-cell responses^31–33^. A major concern is that the glucocorticoids, both synthetic and endogenous, have long been known to drive mice and human DCs towards a tolerogenic state, characterised by impaired upregulation of antigen-presentation and co-stimulation, reduced inflammatory cytokine production and weakened T-cell stimulatory capacity^34,35^. Mechanistically, glucocorticoids signal through the glucocorticoid receptor to transcriptionally reprogram dendritic cells away from full immunogenic maturation and toward a tolerogenic state. Glucocorticoid induced leucine zipper (GILZ) is a central mediator. Its glucocorticoid-induced expression restrains NF-kB and AP-1 inflammatory programs, lowering IL-12, co-stimulatory function, antigen-uptake-linked activation, and T-cell priming. In parallel, glucocorticoid-conditioned DCs acquire broader GR-driven transcriptional and epigenetic remodelling; MAFB stabilizes this tolerogenic programme^36,37^. However, how glucocorticoids remodel chromatin architecture and the epigenomic landscape of GBM-associated DCs, including glucocorticoid receptor (NR3C1) chromatin occupancy patterns remains largely unexplored. These questions are particularly relevant in GBM, where dexamethasone is widely used in clinical practice, yet current DC vaccine approaches have not been developed to resist glucocorticoid-mediated suppression.

Here, we hypothesised that glucocorticoid signalling through NR3C1 is a tumour-intrinsic barrier to DC immunogenicity in GBM, and that disrupting this pathway could unlock the therapeutic potential of DC vaccination. Combining targeted metabolomics and transcriptomic profiling of GBM samples with mechanistic studies in mouse models and patient-derived dendritic cells, we show that glucocorticoids accumulate within the GBM microenvironment and drives NR3C1-dependent chromatin remodelling that suppresses antigen-presentation and co-stimulatory programmes in DCs. Genetic deletion or pharmacological inhibition of NR3C1 reverses this tolerogenic state, restoring DC activation, enhancing cytotoxic CD8+ T-cell priming and improving the efficacy of tumour lysate-loaded DC vaccination *in vivo*. These findings establish glucocorticoid signalling as a defined, targetable mechanism of DC dysfunction in GBM and provide a rationale for steroid-resistant DC vaccines as a strategy to overcome a major clinical barrier to immunotherapy in this disease.

## Results

### Local steroid signalling is associated with impaired dendritic-cell activation in glioblastoma

To define the steroid landscape of GBM and its potential relationship to intratumoural immune dysfunction, we profiled primary human GBM specimens across complementary molecular modalities. We performed targeted LC–MS steroid metabolomics on 24 primary tumours, bulk RNA-seq on 80 tumours, single-nucleus RNA-seq (snRNA-seq) on 12 tumours and integrated these analyses with public single-cell RNA-seq datasets from 28 GBM samples (Fig. 1A). This multi-layered strategy enabled us to jointly assess local steroid abundance, steroid receptor expression and immune-cell state in human GBM. Targeted metabolomics detected multiple endogenous steroids in tumours of GBM patients (Table S1), including pregnenolone, 17-hydroxypregnenolone, dehydroepiandrosterone and cortisol, indicating the presence of an intratumoural steroid-rich milieu (Fig. 1B). Notably, dexamethasone was present at markedly higher levels than endogenous steroids in a subset of tumours, consistent with exogenous synthetic glucocorticoid exposure and highlighting the potential for strong pharmacological engagement of steroid signalling pathways within the tumour microenvironment (Fig. 1B). These findings suggest that both endogenous steroid hormones and clinically administered synthetic glucocorticoids may shape immune states in GBM.

**Fig. 1.**
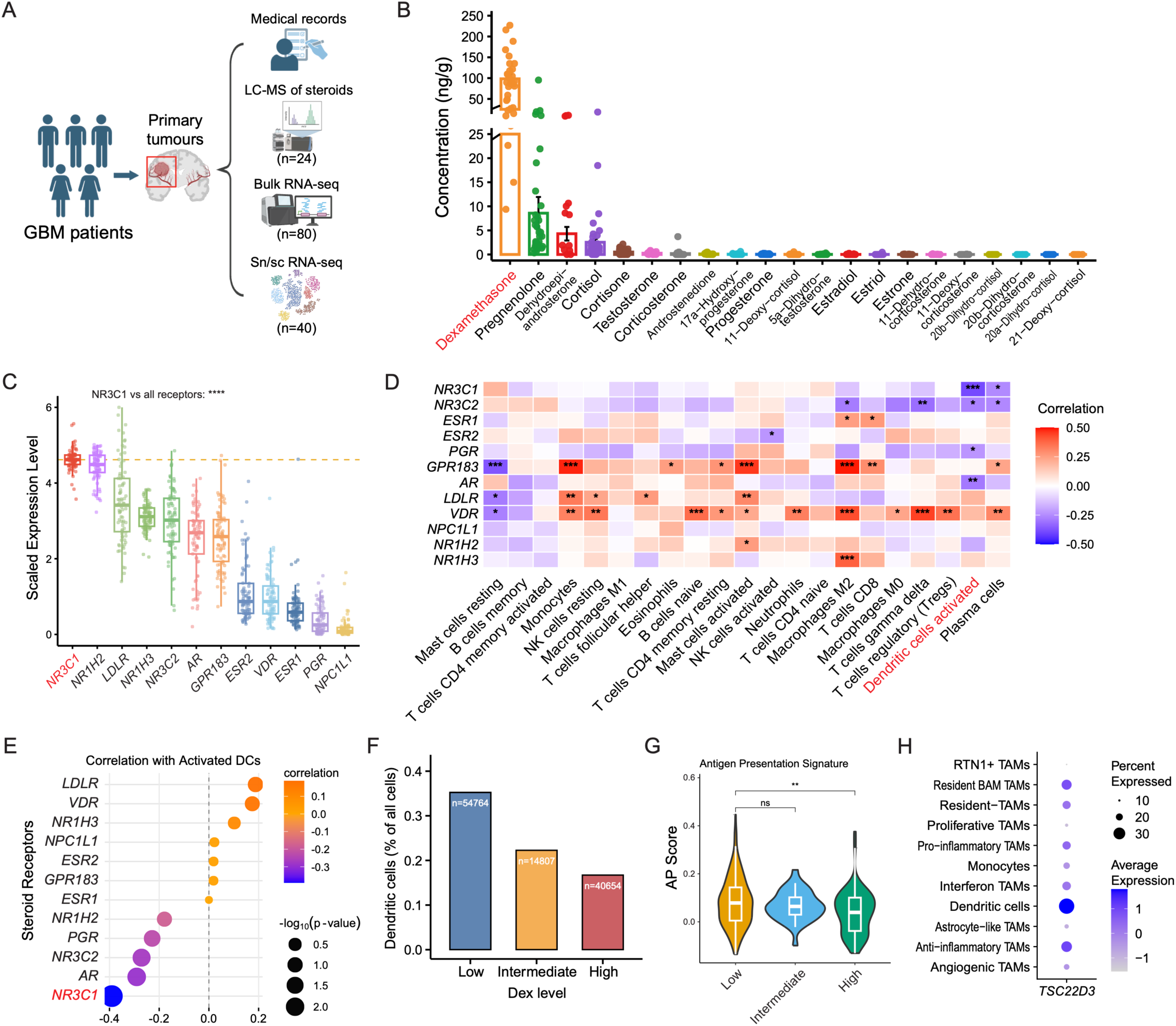
Local steroid signalling is associated with impaired dendritic-cell activation in glioblastoma. **A**, Schematic of the study design. Primary human GBM samples were analysed by quantitative LC–MS/MS steroid metabolomics (n = 24), bulk RNA-seq (n = 80) and sn/scRNA-seq with paired medical-record information (n = 40). **B,** Targeted LC–MS/MS quantification of steroids in primary GBM tumours, showing endogenous steroid hormones together with dexamethasone levels from 24 GBM patients. **C,** Bulk RNA-seq analysis of steroid-receptor expression in GBM tumours, showing the expression of *NR3C1* and other steroid receptors. **D,** Pearson correlation heatmap showing associations between steroid-receptor expression and inferred immune cell abundance quantified by CIBERSORTx across bulk GBM RNA-seq samples. The colour indicates the Pearson correlation coefficient, with red representing positive correlations and blue representing negative correlations. Asterisks indicate statistical significance of Pearson correlation analysis: *P < 0.05, **P < 0.01, ***P < 0.001. **E,** Ranked correlation of steroid-receptor expression with activated DC abundance, highlighting the strongest negative association for *NR3C1*. Dot size indicates significance and colour indicates correlation coefficient. **F,** DC cluster frequence in snRNA-seq data stratified by low, intermediate and high dexamethasone exposure groups defined from paired clinical records. **G,** Antigen-presentation signature scores in DCs across low, intermediate and high dexamethasone exposure groups. **H,** Dot plot showing *TSC22D3* expression across myeloid immune clusters in snRNA-seq data. Dot size represents the percentage of cells expressing *TSC22D3*, and colour indicates average expression level.

We next determined which steroid-sensing pathways are most prominent in GBM. Across bulk RNA-seq data, *NR3C1*, encoding the glucocorticoid receptor, showed the highest expression among major steroid hormone receptors, exceeding androgen receptor (*AR*), oestrogen receptor alpha (*ESR1*), oestrogen receptor beta (*ESR2*), and progesterone receptor (*PGR*) in GBM tumours (Fig. 1C). Consistently, analysis of a public scRNA-seq dataset revealed prominent *NR3C1* expression across GBM-infiltrating immune-cell populations (Figure S1A)^38^. To explore the immunological relevance of this axis, we correlated steroid-receptor expression in bulk transcriptomes with inferred immune-cell abundance using CIBERSORT. Although several receptors showed variable associations across immune populations, *NR3C1* demonstrated a negative association with activated DCs, whereas other steroid receptors displayed weaker or divergent relationships (Fig. 1D, E). Because activated DCs are central to antigen processing, presentation and T-cell priming, this pattern suggested that glucocorticoid signalling may be linked to impaired DC function in GBM.

We also utilised clinical information paired with snRNA-seq data to explore dexamethasone-exposure and DC response. Stratification of samples by low, intermediate and high dexamethasone exposure revealed a progressive reduction in DC abundance with increasing dexamethasone level (Fig. 1F). Consistent with this, DC antigen-presentation signature scores were lower in the high-dexamethasone group than in the low-dexamethasone group (Fig. 1G), supporting the idea that glucocorticoid exposure is associated with functionally restrained DC states in GBM. In parallel, *TSC22D3*, a canonical glucocorticoid-responsive immunoregulatory gene, was expressed higher in DCs than several myeloid clusters (Fig. 1H, S2B). Together, these data establish a clinically relevant association between dexamethasone exposure, NR3C1 signalling and impaired DC abundance and antigen-presenting capacity in human GBM. These findings identified NR3C1 as the leading receptor candidate through which dexamethasone may constrain DC function, prompting us to determine whether dexamethasone directly engages NR3C1 and reprogrammes the regulatory state of patient-derived DCs.

### Dexamethasone engages an NR3C1-centred suppressive regulatory programme in GBM patient-derived dendritic cells

Having identified an association between dexamethasone exposure, NR3C1 and impaired DC function in human GBM, we next asked whether dexamethasone directly engages NR3C1 in patient-derived DCs and whether changes in receptor occupancy are coupled to chromatin and transcriptional reprogramming. We collected peripheral blood samples from 20 patients with GBM, differentiated patient-derived DCs in the presence or absence of dexamethasone, and separately profiled NR3C1 occupancy and chromatin accessibility using CUT&Tag-seq and ATAC-seq. We then integrated these datasets with our previously generated RNA-seq data from glucocorticoid-treated human DCs^19^ (Fig. 2A). This approach enabled us to determine whether dexamethasone directly rewires glucocorticoid receptor binding landscapes in patient-derived DCs and how these regulatory changes relate to transcriptional reprogramming. Genomic annotation of differential NR3C1-bound regions showed that dexamethasone-responsive peaks were distributed across promoter, intronic, downstream, and distal intergenic regions, indicating broad remodelling of both promoter-proximal and distal regulatory elements (Fig. 2B, Table S2). Motif enrichment analysis showed that dexamethasone-increased peaks were enriched for the canonical GRE/NR3C1 motif, supporting ligand-induced, sequence-specific NR3C1 recruitment, together with BACH1/NRF2-like, CHOP/ATF4-like, AP-1 and STAT1 motifs (Fig. 2C). By contrast, dexamethasone-decreased peaks were enriched for PU.1, composite PU.1–IRF8, IRF8, IRF3 and NF-κB motifs (Fig. 2C). Principal component analysis of transcription start site (TSS)-centred NR3C1 CUT&Tag profiles clearly separated control and dexamethasone-treated DCs, demonstrating that glucocorticoid exposure induced a distinct NR3C1 chromatin-occupancy state (Fig. 2D). Differential analysis of NR3C1 occupancy at TSS-centred regions identified marked dexamethasone-associated gains and losses of NR3C1 binding, including previously implicated glucocorticoid-responsive loci, including *FKBP5*, *MAFB* and *TSC22D3*. Importantly, the analysis also identified candidate NR3C1-regulated loci not previously defined in human DCs, including increased occupancy at *C5AR1* and reduced occupancy near *JAML* and *SLC38A2* (Fig. 2E, F and Table S3). Integration with RNA-seq demonstrated concordant increases in NR3C1 occupancy and gene expression at *FKBP5, MAFB, C5AR1, C5AR2* and *TSC22D3*, together with concordant decreases at DC activation-associated loci including *NR4A3, CCR7* and *TRAF1* (Fig. S2A).

**Fig. 2.**
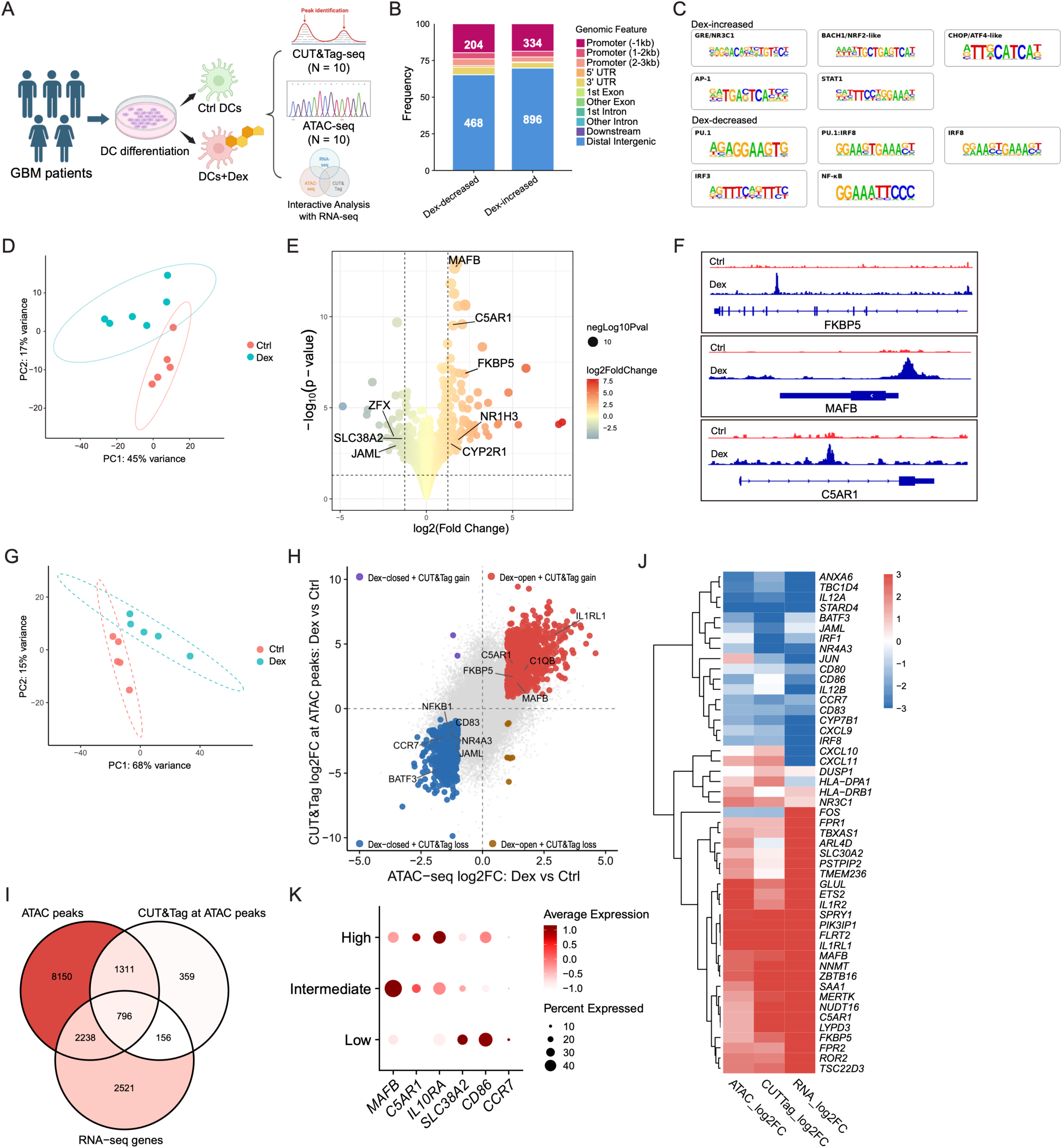
Dexamethasone reprogrammes GBM patient-derived dendritic cells through NR3C1-dependent chromatin remodelling. **A**, Schematic of the experimental design. Peripheral blood samples were collected from patients with glioblastoma, and patient-derived dendritic cells were differentiated in the presence or absence of dexamethasone. Cells were separately profiled by NR3C1 CUT&Tag-seq and ATAC-seq, with integration of RNA-seq data. **B,** Genomic annotation of differential NR3C1 CUT&Tag peaks, stratified by dexamethasone-decreased and dexamethasone-increased regions. Stacked bars indicate the proportion of peaks assigned to promoter, untranslated region (UTR), exon, intron, downstream and distal intergenic regions. **C,** Motif enrichment analysis of differential NR3C1-bound regions from control and dexamethasone-treated DCs. **D,** Principal component analysis of NR3C1 CUT&Tag-seq profiles from control and dexamethasone-treated DCs. **E,** Volcano plot of differential NR3C1 CUT&Tag signal at transcription start site (TSS)-centred regions in dexamethasone-treated versus control DCs. Dot colour indicates log2 fold change and dot size indicates −log10 P value. Selected TSS-associated genes are labelled. **F,** Representative genome-browser tracks showing NR3C1 CUT&Tag signal at selected loci in control and dexamethasone-treated DCs. **G,** Principal component analysis of NR3C1 CUT&Tag signal quantified at ATAC-defined accessible peaks in control and dexamethasone-treated DCs. **H,** Integrated comparison of differential NR3C1 CUT&Tag signal at ATAC peaks and differential chromatin accessibility measured by ATAC-seq in dexamethasone-treated versus control DCs. Selected genes are labelled. **I,** Venn diagram showing overlap among genes or regions identified by ATAC-seq, NR3C1 CUT&Tag at ATAC peaks and RNA-seq analyses. **J,** Heatmap showing integrated ATAC-seq, NR3C1 CUT&Tag and RNA-seq changes for selected overlapping genes. **K,** Dot plot showing expression of selected genes in dendritic cells from snRNA-seq data stratified by low, intermediate and high dexamethasone exposure. Dot size indicates the percentage of cells expressing each gene, and colour indicates average expression level.

ATAC-seq independently revealed a treatment-associated shift in global chromatin accessibility and extensive dexamethasone-induced opening and closing of regulatory regions (Fig. S2B, C). To relate these accessibility changes to receptor binding, we quantified NR3C1 CUT&Tag signals across ATAC-defined peaks. Principal component analysis again showed clear separation between control and dexamethasone-treated samples, indicating widespread remodelling of NR3C1 engagement at accessible regulatory regions (Fig. 2G). Integrated comparison of CUT&Tag signal at ATAC peaks with ATAC-seq changes revealed coordinated glucocorticoid-dependent alterations in both receptor occupancy and chromatin accessibility (Fig. 2H). Among loci showing concordant gain of both accessibility and NR3C1 signal were *FKBP5, C5AR1, MAFB* and *IL1RL1*, whereas loci such as *CCR7, NR4A3, JAML* and *CD83* showed coordinated loss, consistent with reduced DC activation and migratory programmes. Integration of CUT&Tag, ATAC-seq and RNA-seq identified 796 genes shared across the three regulatory layers (Fig. 2I, Table S4). Within this integrated gene set, dexamethasone-associated increases included *FKBP5, TSC22D3, MERTK, NFKB1, IL1R2* and *MAFB*, whereas genes supporting DC activation, migration and antigen-presentation competence, including *CD86, IL12B, CCR7, JAML* and *NR4A3*, were reduced (Fig. 2J). We next examined DCs from the snRNA-seq dataset stratified by documented dexamethasone exposure. *MAFB* and *C5AR1* were preferentially expressed in intermediate- or high-exposure samples, whereas *SLC38A2, CD86* and *CCR7* were enriched in the low-exposure group (Fig. 2K). Together, these data show that dexamethasone directly engages NR3C1 and couples receptor redistribution to chromatin and transcriptional changes that favour glucocorticoid-responsive and dysfunctional DC related gene expression, prompting us to test whether selective deletion of *Nr3c1* in DCs could overcome dexamethasone-associated immune dysfunction *in vivo*.

### DC-specific deletion of *Nr3c1* restricts glioblastoma growth and restores anti-tumour immunity *in vivo*

To test whether glucocorticoid signalling in DCs functionally constrains anti-tumour immunity in GBM, we generated DC-specific *Nr3c1* conditional knockout mice by crossing *Nr3c1*^fl/fl^ mice with *Cd11c*^Cre^ mice. We implanted SB28 GBM cells and treated tumour-bearing mice in the presence or absence of dexamethasone to model the glucocorticoid-exposed clinical setting. Tumour progression was monitored longitudinally, and tumour-infiltrating CD45^+^ immune cells were isolated for immunpphenotyping by scRNA-seq and flow-cytometry (Fig. 3A). *Cd11c*^Cre^ deletes *Nr3c1* very efficiently in *Nr3c1*^fl/fl^;*Cd11c*^Cre^ (Nr3c1 cKO) mice (Fig. 3B). Under dexamethasone exposure, DC-specific deletion of *Nr3c1* restrained tumour progression, with reduced tumour size, slower tumour growth and lower endpoint tumour weight compared with control mice (Fig. 3C–E). By contrast, in the absence of dexamethasone, DC-specific *Nr3c1* deletion produced only a modest reduction in tumour growth (Fig. S3A, B). These findings indicate that the anti-tumour effect of DC-specific *Nr3c1* deletion is most evident under glucocorticoid pressure *in vivo*, supporting a specific role for DC-intrinsic *NR3C1* in dexamethasone-associated immune suppression.

**Fig. 3.**
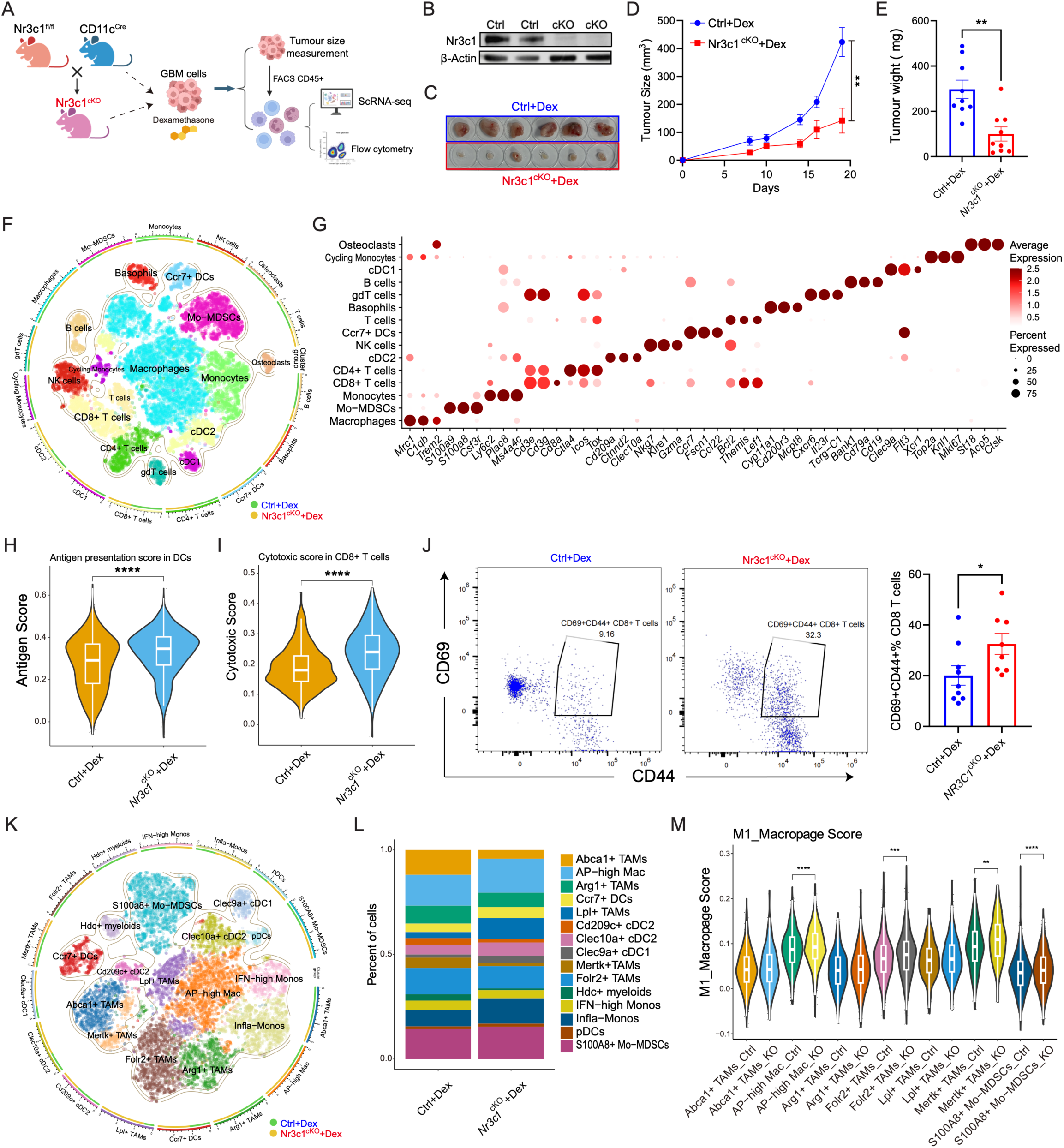
Dendritic cell-specific deletion of Nr3c1 restricts glioblastoma growth and restores anti-tumour immunity in vivo. **A**, Schematic of the *in vivo* experimental design. DC-specific *Nr3c1* conditional knock out (*Nr3c*1^fl/fl^;*Cd11c*^Cre^ or *Nr3c1*^cKO^) and control mice (*Cd11c*^Cre^) were implanted with SB28 glioblastoma cells under dexamethasone treatment, followed by tumour-size monitoring, isolation of tumour-infiltrating CD45^+^ immune cells, scRNA-seq and flow-cytometric analysis. **B,** Western blot showing Nr3c1 expression in control (Nr3c1-sufficient) and *Nr3c1* conditional knockout (*Nr3c*1^fl/fl^;*Cd11c*^Cre^) bone marrow-derived DCs. **C,** Representative tumour images from control and *Nr3c1*^cKO^ (*Nr3c*1^fl/fl^;*Cd11c*^Cre^) mice under dexamethasone treatment. **D,** Tumour growth curves in control and *Nr3c1*^cKO^ mice under dexamethasone treatment. Data are presented as mean ± SEM. Statistical significance was assessed using a two-way mixed-effects ANOVA model with treatment group and time as factors. *P < 0.05, **P < 0.01, ***P < 0.001. **E,** Endpoint tumour weight in control and *Nr3c1*^cKO^ mice under dexamethasone treatment. **F,** UMAP projection of scRNA-seq profiles from tumour-infiltrating CD45^+^ immune cells to show the differences in major immune cell composition (immunophenotyping). coloured by annotated immune-cell populations. **G,** Dot plot showing expression of selected marker genes for specific tumour-infiltrating immune-cell populations. Dot size indicates the percentage of cells expressing each gene, and colour indicates average expression level. **H,** Antigen-presentation signature scores in tumour-infiltrating DCs from control and *Nr3c1*^cKO^ mice as observed in scRNA-seq data obtained from tumour-infiltrating CD45^+^ immune cells **I,** Analysis of comparative cytotoxicity scores in tumour-infiltrating single-cell transcriptome of CD8^+^ T cells from control and *Nr3c1*^cKO^ mice. **J,** Representative flow cytometry plots and quantification of CD69^+^CD44^+^ CD8^+^ T cells in tumours from control and *Nr3c1*^cKO^ knockout mice. **K,** UMAP projection of re-clustered tumour-infiltrating myeloid cells, coloured by annotated macrophage, monocyte and dendritic-cell states. **L,** Stacked bar plot showing the relative abundance of annotated myeloid-cell populations in control and *Nr3c1*^cKO^ tumours. **M,** M1 macrophage signature scores across annotated myeloid-cell populations in control and *Nr3c1*^cKO^ tumours.

To define the immunological basis of this phenotype, we profiled tumour-infiltrating immune cells by scRNA-seq. Global embedding resolved diverse lymphoid and myeloid populations, including T, B and NK cells, monocytes, macrophages, Mo-MDSCs and multiple DC subsets, including cDC1, cDC2 and CCR7^+^ DCs (Fig. 3F). These annotations were supported by canonical lineage markers, including *Cd3d/Cd3e* for T cells, *Nkg7* for NK cells, *Ms4a1/Cd79a* for B cells, *Ccr7/Fscn1* for migratory DCs, *Clec9a/Xcr1* for cDC1 and *Cd209a/Clec10a* for cDC2 (Fig. 3G). *Nr3c1*-deficient tumour-infiltrating DCs displayed higher antigen-presentation scores than control DCs (Fig. 3H). Because enhanced DC antigen presentation is expected to influence downstream T-cell immunity, we next examined tumour-infiltrating T cells. Re-clustering resolved seven transcriptionally distinct states, including naïve T cells, central memory T cells, cytotoxic CD8^+^ T cells, conventional CD4^+^ T cells, regulatory T cells, γδT17 cells and IFN-high T cells (Fig. S3C, D). Consistent with enhanced DC antigen presentation, tumour-infiltrating CD8⁺ T cells from *Nr3c1* cKO mice exhibited higher cytotoxicity signature scores than those from control mice (Fig. 3I). We also observed an increased frequency of activated CD69^+^CD44^+^ CD8^+^ T cells in the knockout group (Fig. 3J), supporting enhanced T-cell activation *in vivo*. We next investigated whether DC-specific *Nr3c1* deletion also remodelled the broader myeloid compartment. Re-clustering explored 15 myeloid states, including Abca1^+^, Mertk^+^, Lpl^+^, Folr2^+^ and Arg1^+^ TAM populations, antigen-presentation-high macrophages, inflammatory and IFN-high monocytes, S100A8^+^ Mo-MDSCs, and multiple DC subsets (Fig. 3K, S3F). Quantification of myeloid-cell composition showed a reduction in tumour-associated macrophage states, including Abca1^+^ and Mertk^+^ TAM-like populations, alongside changes in inflammatory monocyte and DC subsets (Fig. 3L). Consistent with a less suppressive myeloid landscape, M1-like macrophage scores were increased in many myeloid populations in the knockout tumours (Fig. 3M). Together, these findings show that selective disruption of glucocorticoid receptor signalling in DCs restores antigen-presenting function, enhances CD8^+^ T-cell activation and remodels the GBM myeloid microenvironment towards a more inflammatory anti-tumour state under dexamethasone exposure.

### Genetic deletion of *NR3C1* confers steroid resistance and restores functional competence in patient-derived dendritic cells

To determine whether the immunosuppressive effects of glucocorticoids in human DCs can be reversed therapeutically or by genetic engineering, we established two complementary approaches in GBM patient-derived DCs: pharmacological glucocorticoid receptor blockade using mifepristone (Mifeprex) and non-viral CRISPR–Cas9-mediated deletion of *NR3C1* (Fig. 4A). Flow-cytometric analysis showed that dexamethasone treatment reduced the expression of the co-stimulatory molecule CD86, whereas mifepristone (Mifeprex) restored CD86 expression when cultured with dexamethasone (Fig. 4B). CRISPR editing is a viable strategy to achieve stable disruption of glucocorticoid signalling. We reasoned that the resultant DC could be suitable to improve the utility of DC vaccine in GBM patient. We therefore developed guide RNAs targeting *NR3C1* using non-viral CRISPR–Cas9 ribonucleoprotein delivery. Delivery of NR3C1-targeting guide RNAs after DC differentiation but before maturation yielded the highest editing efficiency and was therefore used for downstream functional studies. Sanger sequencing and indel analysis confirmed efficient *NR3C1* disruption, with selected editing conditions and gRNAs achieving high knockout efficiency in patient-derived DCs (Fig. 4C). Functionally, NR3C1-deficient DCs maintained higher CD86 expression despite dexamethasone exposure compared with Cas9-control DCs, supporting restoration of co-stimulatory competence after genetic receptor ablation (Fig. 4D).

**Fig. 4.**
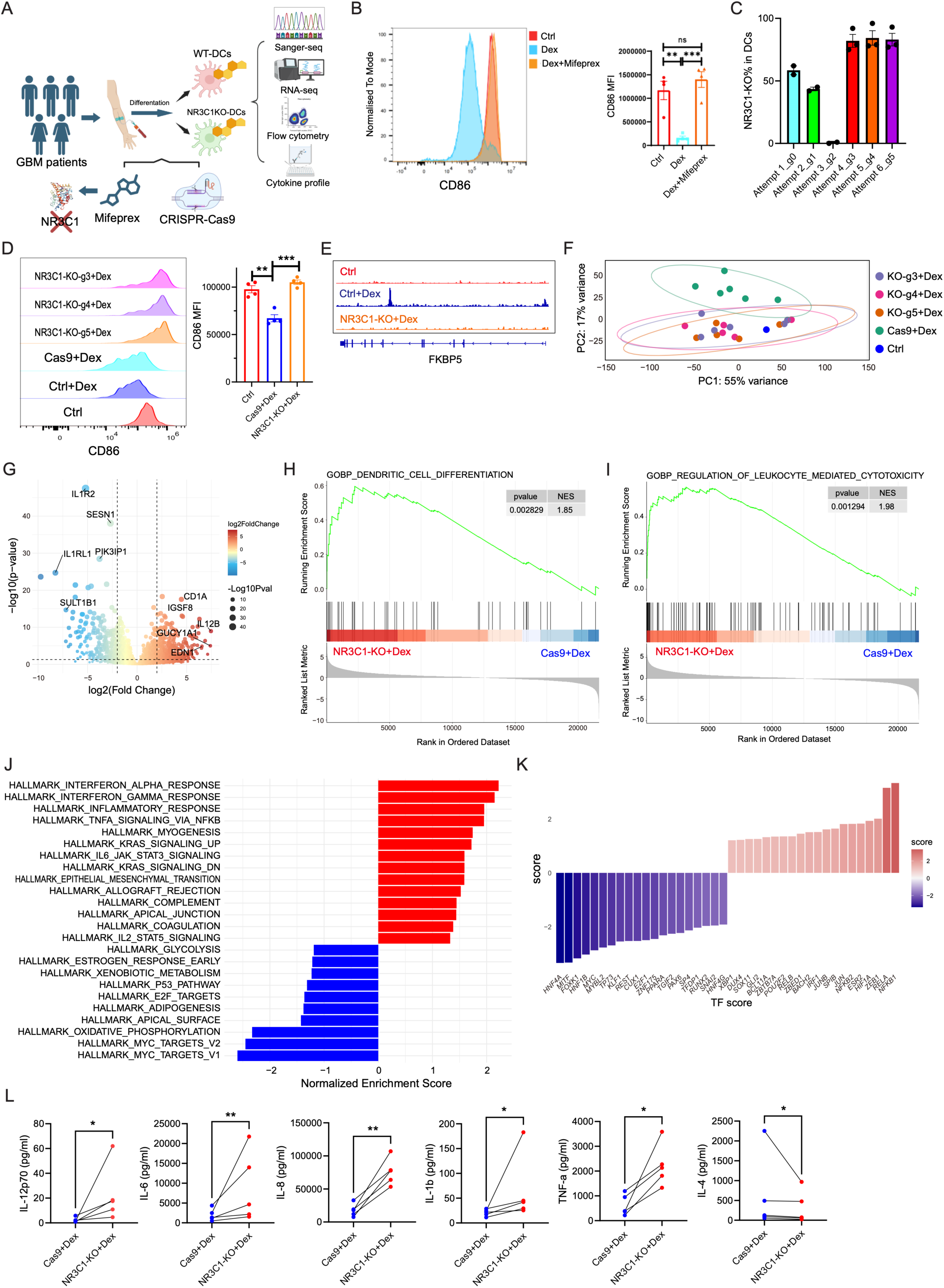
NR3C1 deletion confers steroid resistance and restores dendritic-cell function in GBM patient-derived DCs. **A**, Schematic of the experimental strategy. Peripheral blood samples were collected from patients with glioblastoma, monocyte-derived DCs were generated *ex vivo*, and cells were subjected either to pharmacological glucocorticoid receptor blockade with mifepristone or to non-viral CRISPR–Cas9-mediated *NR3C1* disruption. Downstream analyses included flow cytometry, CUT&Tag-seq, RNA-seq and cytokine profiling. **B,** Representative flow-cytometry histogram (left) and quantification (right) showing CD86 expression in control (dexamethasone-untreated and *NR3C1* sufficient), dexamethasone-treated and mifepristone-treated DCs. **C,** Indel frequencies for NR3C1-targeting guide RNAs assessed by Sanger sequencing/ICE analysis to show the knockout efficiency. **D,** Representative flow-cytometry histograms and quantification of CD86 expression in Cas9-control and NR3C1-edited DCs under the indicated treatment conditions. **E,** Representative genome-browser tracks showing NR3C1 CUT&Tag signal at the FKBP5 locus in control, Cas9-control and NR3C1-edited DCs under the indicated conditions. **F,** Principal component analysis of RNA-seq profiles from Cas9-control and NR3C1-edited DCs under control or dexamethasone-treated conditions. **G,** Volcano plot showing differentially expressed genes in *NR3C1*-edited versus Cas9-control DCs under dexamethasone exposure. Dot colour indicates log2 fold change and dot size indicates −log10 P value. Selected genes are labelled. **H,I**, Gene set enrichment analysis (GSEA) plots showing enrichment of representative pathways (DC differentiation and regulation of leukocyte mediated cytotoxicity) in *NR3C1*-edited versus Cas9-control DCs under dexamethasone exposure. **J,** Hallmark pathway enrichment analysis showing normalised enrichment scores for pathways differentially represented in *NR3C1*-edited versus Cas9-control DCs. **K,** Transcription-factor activity analysis showing differentially enriched transcription-factor programmes in NR3C1-edited versus Cas9-control DCs. **L,** Cytokine profiling of culture supernatants from Cas9-control and *NR3C1*-edited DCs under dexamethasone exposure, showing selected cytokines measured by multiplex assay.

We next examined whether NR3C1 deletion restored DC activation at the molecular and functional levels using CUT&Tag-seq, RNA-seq, and cytokine profile. NR3C1 CUT&Tag tracks at the canonical glucocorticoid-responsive FKBP5 locus confirmed dexamethasone-induced NR3C1 chromatin occupancy in Cas9-control DCs and loss of this signal following *NR3C1* knockout (Fig. 4E). RNA-seq further showed that NR3C1-deficient DCs occupied a transcriptional state distinct from dexamethasone-treated Cas9-control DCs (Fig. 4F), indicating broad molecular reprogramming after glucocorticoid receptor ablation. Differential expression analysis identified substantial transcriptional changes in NR3C1-deficient DCs under dexamethasone exposure (Fig. 4G, Table S5). Upregulated genes included CD1A, IL12B, IGSF8, GUCY1A1 and EDN1, consistent with enhanced antigen-presenting and inflammatory programmes. Conversely, genes associated with glucocorticoid-responsive or restrained DC states, including IL1R2, IL1RL1, SESN1 and PIK3IP1, were reduced in NR3C1-deficient DCs (Fig. 4G). To determine whether *NR3C1* deletion specifically reversed the regulatory programme imposed by dexamethasone, we integrated dexamethasone-regulated ATAC-seq, NR3C1 CUT&Tag and RNA-seq datasets with RNA-seq changes following *NR3C1* knockout. This analysis identified 198 genes altered across chromatin accessibility, NR3C1 occupancy, dexamethasone-induced expression and NR3C1-knockout-associated expression (Fig. S4A). Among these, dexamethasone-induced genes, including *FKBP5, TSC22D3, IL1RL1, PIK3IP1, C5AR1* and *MAFB*, were reduced following *NR3C1* deletion, whereas dexamethasone-repressed DC activation genes, including *CD80, CD86, IL12A, IL12B, CCR7, CD83, BATF3* and *JAML*, were restored (Fig. S4B).

Gene set enrichment analysis further showed enrichment of pathways related to DC differentiation, regulation of leukocyte-mediated cytotoxicity, antigen processing and presentation of endogenous antigens, and inflammatory responses to antigenic stimuli in NR3C1-deficient DCs (Fig. 4H-I, S4C), supporting restoration of DC programmes required for T-cell stimulation. At the broader pathway level, *NR3C1* deletion enhanced inflammatory and immune-activating signatures, including interferon, TNF–NF-κB, and cytokine-associated programmes, while reducing stress-associated pathways (Fig. 4J). Transcription-factor activity analysis revealed widespread regulatory rewiring in NR3C1-deficient DCs, consistent with release from dexamethasone-imposed transcriptional restraint (Fig. 4K). To determine whether *NR3C1* deletion restores DC secretory function under glucocorticoid exposure, we profiled culture supernatants from Cas9-control and *NR3C1*-edited DCs treated with dexamethasone using a multiplex cytokine assay. *NR3C1*-KO DCs secreted significantly higher levels of inflammatory and T-cell-supportive cytokines IL-12p70, IL-6, IL-8, IL-1β, and TNF-α compared with Cas9-control DCs under dexamethasone exposure (Fig. 4L). In contrast, IL-4 secretion was significantly reduced in *NR3C1*-KO DCs relative to Cas9-control DCs under the same conditions (Fig. 4L). Together, these data show that non-viral CRISPR–Cas9-mediated *NR3C1* deletion confers steroid resistance and restores functional competence in GBM patient-derived dendritic cells making them appropriate for DC vaccine.

### *Nr3c1*-deficient dendritic cells enhance antigen-specific CD8^+^ T-cell responses and improve glioblastoma control *in vivo*

To determine whether the functional recovery of *Nr3c1*-deficient DCs translates into improved antigen-specific T-cell priming, we generated bone marrow-derived DCs (BMDCs) from control and *Nr3c1* conditional knockout mice, pulsed them with OVA peptide in the presence or absence of dexamethasone, and co-cultured them with CellTrace Violet labelled CD8^+^ T cells isolated from OT-I mice (Fig. 5A). CellTrace Violet dilution analysis showed that OT-I CD8^+^ T cells underwent greater proliferation when primed with dexamethasone-exposed *Nr3c1*-deficient DCs than when primed with control DCs from wildtype mice (Fig. 5B). Consistent with enhanced activation as revealed from enhanced proliferation, flow-cytometric analysis further demonstrated an increased frequency of CD69^+^CD44^+^ CD8^+^ T cells in cultures primed with *Nr3c1*-deficient DCs (Fig. 5C). These data indicate that deletion of *Nr3c1* restores the capacity of DCs to drive productive antigen-specific CD8^+^ T-cell responses despite glucocorticoid exposure.

**Fig. 5.**
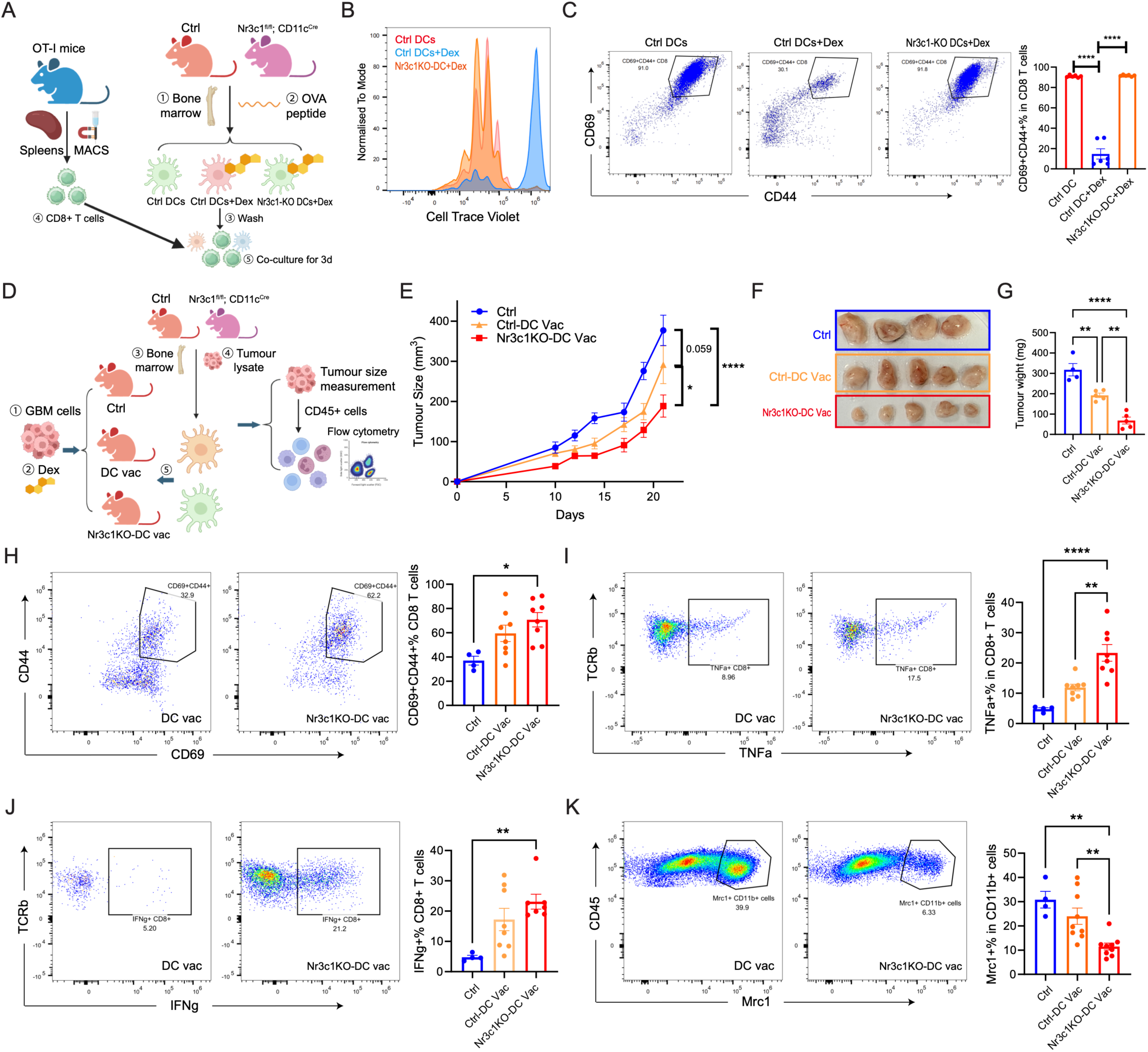
Nr3c1-deficient dendritic cells enhance antigen-specific CD8^+^ T-cell responses and improve glioblastoma control *in vivo*. **A**, Schematic of the *in vitro* antigen-presentation assay. Bone marrow-derived dendritic cells (BMDCs) from control or DC-specific *Nr3c1* knockout mice were pulsed with OVA peptide under the indicated conditions and co-cultured with splenic CD8^+^ T cells isolated from OT-I mice. **B,** Representative CellTrace Violet dilution histograms showing proliferation of OT-I CD8^+^ T cells after co-culture with control or Ova-pulsed Nr3c1-deficient DCs. **C,** Representative flow-cytometry plots and quantification of CD69 and CD44 expression on OT-I CD8^+^ T cells after co-culture with control DCs, dexamethasone-treated control DCs or dexamethasone-treated Nr3c1-deficient DCs. **D,** Schematic of the *in vivo* dendritic cell therapeutic experiment. SB28 tumour-bearing mice under dexamethasone treatment received tumour lysate-loaded Nr3c1-sufficient (control) or Nr3c1-deficient DC vaccines, followed by tumour monitoring and immune profiling. **E,** Tumour growth curves for untreated control mice, mice treated with control DC vaccines and mice treated with Nr3c1-deficient DC vaccines. Error bars are presented as mean ± SEM. Statistical significance was assessed using a two-way mixed-effects ANOVA model with treatment group and time as factors. *P < 0.05, **P < 0.01, ***P < 0.001. **F,** Representative images of excised tumours from the indicated treatment groups at endpoint. **G,** Endpoint tumour weight in the indicated treatment groups. **H,** Representative flow-cytometry plots and quantification of CD69^+^CD44^+^ CD8^+^ T cells in tumours from the indicated treatment groups. **I,** Representative flow-cytometry plots (left-panel) and quantification (right-panel) of TNFα^+^ CD8^+^ T cells in tumours from the indicated treatment groups. **J,** Representative flow-cytometry plots (left-panel) and quantification (right-panel) of IFNγ^+^ CD8^+^ T cells in tumours from the indicated treatment groups. **K,** Representative flow-cytometry plots and quantification of Mrc1^+^ CD11b^+^ cells in tumours from the indicated treatment groups.

We next tested whether this enhanced T-cell priming capacity could improve anti-tumour efficacy *in vivo*. SB28 tumour lysate-loaded control (Nr3c1-sufficient) or Nr3c1-deficient DC vaccines were evaluated in a therapeutic SB28 glioblastoma model under dexamethasone treatment (Fig. 5D). Mice receiving Nr3c1-deficient DC vaccines (i.e., adoptive transfer of tumour lysate-loaded DC) showed improved tumour control, with reduced tumour growth over time, smaller excised tumours and lower tumour weight at endpoint compared with mice treated with control DC vaccines (Nr3c1-sufficient) or untreated (without DC vac) controls (Fig. 5E–G). To define the accompanying immune changes, we analysed tumour-infiltrating T-cell responses by flow cytometry. Treatment with Nr3c1-deficient DC vaccines increased the frequency of activated CD69^+^CD44^+^ CD8^+^ T cells (Fig. 5H), together with higher proportions of TNFα^+^ and IFNγ^+^ CD8^+^ T cells (Fig. 5I, J), suggesting an enhanced effector immunity downstream of improved antigen presentation and priming. In parallel, tumours from mice treated with Nr3c1-deficient DC vaccines contained fewer Mrc1^+^CD11b^+^ myeloid cells than control groups (Fig. 5K), suggesting a reduction in immunosuppressive myeloid features within the tumour microenvironment. Together, these data show that *Nr3c1* deletion provides DCs with stronger T-cell-stimulatory capacity and enhances the therapeutic efficacy of lysate-loaded DC vaccination in mouse model of SB28 glioblastoma.

## Discussion

Our study identifies glucocorticoid signalling as a targetable mechanism of dendritic-cell dysfunction in glioblastoma. By integrating steroid metabolomics, bulk and single-cell immune profiling, patient-derived DC perturbation, NR3C1 CUT&Tag/ATAC-seq, mouse genetics and functional DC-vaccine assays, we link a steroid-rich GBM microenvironment to impaired DC activation and show that inhibition or deletion of glucocorticoids receptor restores DC function. This multi-layered evidence places NR3C1 upstream of defective antigen presentation, impaired co-stimulation and weakened CD8^+^ T-cell priming, while also connecting DC steroid resistance to broader remodelling of the myeloid tumour microenvironment. Importantly, the work moves beyond treating dexamethasone exposure as a clinical confounder and instead nominates glucocorticoid resistance as an actionable design principle for next-generation DC vaccines in GBM.

Dexamethasone presents a clinical paradox in GBM. It remains one of the most effective and widely used agents for controlling peritumoural oedema and neurological symptoms, and therefore cannot simply be withdrawn from the therapeutic regimen^27,39,40^. However, its immunological cost is increasingly difficult to ignore. Retrospective clinical analyses, preclinical glioma models and recent meta-analyses have associated corticosteroid exposure with poorer survival and a more immunosuppressed glioma microenvironment^28,41–43^. This concern is particularly relevant for immune therapies that depend on intact antigen presentation and T-cell priming. Checkpoint blockade, neoantigen vaccination and DC vaccination may all be vulnerable to steroid pressure, as suggested by reduced benefit from PD-1/PD-L1 blockade in dexamethasone-exposed GBM cohorts and by the impaired vaccine-induced T-cell responses observed in dexamethasone-treated patients in the 2019 GBM neoantigen vaccine study^29,43^. DC vaccines are especially exposed to this biology because their therapeutic mechanism depends directly on DC maturation, co-stimulation, cytokine production and productive CD8^+^ T-cell activation, processes known to be suppressed by glucocorticoids and^44,45^. Our findings provide a mechanistic and translational solution to this problem: rather than relying solely on dexamethasone avoidance, *NR3C1* disruption delivers DCs intrinsically resistant to glucocorticoid-mediated suppression, restores antigen-presentation and inflammatory programmes, enhances CD8^+^ T-cell priming and improves tumour lysate-loaded DC therapy *in vivo*. Thus, steroid-resistant DC engineering may convert dexamethasone from an unavoidable clinical confounder into a design constraint that can be addressed during vaccine manufacture.

Our study also extends the mechanistic understanding of glucocorticoid-induced DC dysfunction by defining its chromatin basis in GBM patient-derived DCs. Although the tools, ATAC-seq and CUT&Tag-seq, have actively improved the chromatin accessibility and transcription-factor occupancy approaches, their application to human DC biology, and especially to GBM-associated DC therapeutics, remains limited^46–48^. Our findings are consistent with prior studies that glucocorticoids can induce tolerogenic DC states through canonical response genes and transcriptional regulators, including FKBP5, TSC22D3/GILZ and cooperative NR3C1–MAFB activity^36,37,49^. Integration of ATAC-seq, NR3C1 CUT&Tag-seq and RNA-seq identified concordant dexamethasone-associated gains in NR3C1 occupancy and chromatin accessibility at loci including *FKBP5, MAFB* and *C5AR1*, together with transcriptional induction of glucocorticoid-responsive genes, whereas reduced occupancy and accessibility at *JAML, CCR7* and *NR4A3* were accompanied by decreased expression of genes associated with DC activation and migration. Alongside established glucocorticoid-responsive genes, we identified additional candidate NR3C1-associated loci, including increased occupancy near *C5AR1* and reduced occupancy near *JAML* and *SLC38A2*. These genes are biologically plausible regulators of DC function: C5a–C5aR1 signalling can promote tolerogenic cancer-associated DC migration ^50^, JAML supports trans-endothelial migration of DC vaccine in cancer immunotherapy ^51^, and SLC38A2-dependent glutamine signalling has recently been shown to tune cDC1-mediated anti-tumour immunity ^52^. Thus, our epigenomic data nominate a set of GBM-relevant glucocorticoids receptor-linked regulatory nodes that may explain how glucocorticoids suppress antigen presentation, co-stimulation and cytokine production in patient-derived DCs, providing a framework for future functional validation and for designing steroid-resistant DC vaccine products.

Interestingly, the *in vivo* genetic data suggest that DC-intrinsic glucocorticoid signalling acts upstream of broader lymphoid and myeloid remodelling in GBM. In Nr3c1 conditional knockout tumours, DCs showed increased antigen-presentation scores, providing a plausible mechanism for the enhanced CD69⁺CD44⁺ CD8⁺ T-cell activation and cytotoxic transcriptional programmes. This interpretation is consistent with extensive evidence that tumour-infiltrating DCs, particularly cross-presenting cDC populations, are required for tumour-antigen transport, CD8⁺ T-cell priming, effector T-cell recruitment and IL-12-dependent anti-tumour immunity^53–56^. Interestingly, the immune remodelling was not restricted to lymphocytes. *Nr3c1* deletion in DCs was also associated with reduced Abca1⁺ and Mertk⁺ TAM-like populations, alongside increased inflammatory monocytes and higher M1-like macrophage scores. GBM is highly enriched in macrophages and microglia, and these cells can support tumour progression, immune suppression and therapy resistance^57–59^. The reduction in Abca1/Mertk-linked TAM features is particularly relevant because lipid recycling, cholesterol efflux and MerTK-dependent efferocytosis have been implicated in tumour-promoting macrophage states, including lipid-laden macrophages that recycle myelin-derived lipids to fuel mesenchymal-like GBM malignancy^60–62^. Together, these findings support a model in which loss of glucocorticoid signalling in DCs restores antigen presentation output, thereby strengthening CD8⁺ T-cell pressure and disrupting a lipid-conditioned, efferocytic TAM niche.

From a translational perspective, our data support NR3C1 inhibition as a practical route to steroid-resistant DC vaccine products. Pharmacological glucocorticoid-receptor blockade with mifepristone showed partial reversibility, but systemic GR antagonism in patients with GBM may be difficult to deploy alongside the clinical need for dexamethasone. By contrast, ex vivo, non-viral CRISPR–Cas9 ribonucleoprotein editing provides a product-centred strategy that the vaccine cells can be engineered during manufacture without viral vectors, permanent vector integration or long-term systemic GR blockade^63,64^. Non-viral CRISPR platforms have already shown feasibility for clinical-grade immune-cell engineering, including multiplex edited T cells and personalised TCR replacement, supporting the broader translational maturity of this approach^65–68^. In our study, CRISPR–Cas9 ribonucleoprotein editing of *NR3C1* achieved high knockout efficiency in patient-derived DCs and restored co-stimulatory, inflammatory, antigen-presentation and CD8^+^ T-cell-regulatory programmes. Importantly, this molecular rescue translated into function that Nr3c1-deficient DCs enhanced antigen-specific CD8^+^ T-cell expansion and activation in the OT-I system, and tumour lysate-loaded Nr3c1-deficient DCs reduced SB28 glioblastoma growth while increasing CD8^+^ T-cell responses and decreasing immunosuppressive TAMs. This is particularly relevant because tumour lysate-loaded DC vaccines have shown clinical promise in GBM but remain heterogeneous in efficacy^14,69^. Thus, rather than viewing dexamethasone exposure only as a patient-level exclusion or stratification variable, our findings suggest that glucocorticoid resistance can be engineered directly into the DC vaccine product.

Nevertheless, further steps will be required before NR3C1-edited steroid-resistant DCs can be advanced towards clinical testing. First, the therapeutic effect should be validated in more clinically faithful GBM settings, including standard-of-care combinations and defined dexamethasone-exposure schedules, because steroid dose, timing and neurological indication are likely to influence both immune fitness and trial eligibility^3,14,70^. Second, the product itself will need to be converted into a reproducible GMP-compatible autologous manufacturing process, including leukapheresis or blood collection, CD14^+^ monocyte enrichment, moDC differentiation, autologous tumour-lysate preparation, non-viral NR3C1 editing, maturation, cryopreservation and release testing^71–73^. Third, potency assays should be prospectively defined to reflect the proposed mechanism of action, rather than relying only on phenotype; for this product, a rational release matrix could include *NR3C1* editing efficiency, viability, sterility, endotoxin and mycoplasma testing, CD80/CD86/HLA-DR expression, IL-12 and inflammatory cytokine production, antigen-presentation signatures, dexamethasone-resistance assays and autologous T-cell priming under steroid challenge^74,75^. Fourth, because this is a genome-edited cellular product, clinical translation will require a formal safety package addressing on-target indel spectrum, off-target editing, structural variants, chromosomal abnormalities, residual Cas9/sgRNA, product persistence, cytokine-release risk and tumour-promoting potential, in line with current genome-editing and genetically modified cell-therapy guidance^76,77^. Finally, early-phase trials should be designed primarily around feasibility, safety and pharmacodynamic immune endpoints, while stratifying patients by dexamethasone exposure and incorporating paired tissue, blood and vaccine-product biomarkers. If these steps are met, steroid-resistant DC engineering could provide a practical way to preserve DC-vaccine function in the many patients with GBM who cannot safely discontinue dexamethasone.

## Methods

### Human specimens and ethical approval

Primary tumour tissue and peripheral blood were obtained from patients with histologically confirmed glioblastoma who were operated at Addenbrooke’s Hospital, Cambridge University Hospitals NHS Foundation Trust. Written informed consent was obtained from all participants before sample collection. The study was approved by the East of England, Cambridge Central Research Ethics Committee, REC no. 23/EE/0241, and was conducted in accordance with the Declaration of Helsinki and local institutional guidelines.

Fresh tumour tissue was divided for metabolomic (steroid profiling) and sequencing analyses where sufficient material was available. Tumour specimens for metabolomics were snap-frozen in liquid nitrogen immediately after surgical resection and stored at −80 °C until extraction.

For blood-based assays, peripheral blood was collected into EDTA or heparinised tubes and processed on the day of collection. Clinical metadata, including dexamethasone exposure at the time of surgery or blood draw, were abstracted from the medical record where available.

### Murine studies and ethical approval

All mice were handled and maintained in accordance with the UK Animals in Science Regulation Unit Code of Practice for the Housing and Care of Animals Bred, Supplied or Used for Scientific Purposes and the Animals (Scientific Procedures) Act 1986 Amendment Regulations 2012. All procedures were performed under UK Home Office Project Licence PPL PP4938782 and were approved by the local Animal Welfare and Ethical Review Body. Sample sizes were determined based on previous experimental experience and a priori power analysis using G*Power. Mice were housed at the Gurdon Animal Facility under specific pathogen-free conditions with a 12-h light/12-h dark cycle. Genotyping was performed by Transnetyx. DC-specific Nr3c1 conditional knockout mice were generated by crossing *Nr3c1*^fl/fl^ mice^78^ with *Itgax*^Cre^ mice ^79^ (also referred to as *Cd11c*^Cre^ mice), from The Jackson Laboratory. Experimental mice were 8–12 weeks old. C57BL/6J wild-type mice were used for adoptive transfer experiments.

### Targeted LC–MS/MS analysis of steroid hormones and dexamethasone in GBM tissues

Targeted steroid and dexamethasone quantification was performed on snap-frozen GBM tissues. Approximately 50 mg frozen tissue was weighed and transferred into 2-ml reinforced tubes containing 1.4-mm ceramic beads (Fisher Scientific). Samples were extracted in 1 ml acetonitrile containing 0.1% formic acid and spiked with 20 μl isotopically labelled steroid internal standard mixture. Tissues were homogenised using a Bead Ruptor 24 Elite fitted with a CryoCool unit (Omni International) at 1 m s⁻¹ for 30 s, for three cycles.

Homogenates were clarified using a Filter^+^ plate (Biotage), and eluates were further processed through a phospholipid depletion plate (PLD^+^; Biotage). Extracts were dried and reconstituted in water:methanol (70:30, v/v) before LC–MS/MS analysis. Liquid chromatography was performed using an I-Class UPLC system (Waters) with a Kinetex C18 column (150 × 2.1 mm, 2.6 μm). The flow rate was 0.3 ml min⁻¹, using water and methanol mobile phases containing 0.05 mM ammonium fluoride. The gradient started at 50% methanol, increased to 95% methanol, and returned to 50% methanol. The column temperature was maintained at 50 °C, the autosampler at 10 °C, and the injection volume was 20 μl. The total run time was 16 min per sample. Steroids and dexamethasone were detected using a QTrap 6500+ mass spectrometer (AB Sciex) equipped with an electrospray ionisation Turbo V ion spray source. Data were acquired in multiple reaction monitoring mode using optimised positive and negative ionisation settings. Positive and negative ion spray voltages were set to 5,500 V and −4,500 V, respectively, with a source temperature of 600 °C. A panel of 18 steroids was separated and quantified. Representative transitions included pregnenolone at m/z 317.1 → 281.1/159.0 and its internal standard ^13^C₂,d₂-pregnenolone at m/z 321.2 → 285.2.

Peak integration and quantification were performed using MultiQuant v3.0.3 (AB Sciex). Analyte-to-internal-standard peak-area ratios were calculated for each steroid, and concentrations were derived by linear regression against calibration standards. Quantified analytes included pregnenolone, aldosterone, progesterone, 17β-estradiol, 5α-dihydrotestosterone, testosterone and dexamethasone. Steroid abundance was normalised to the recorded tissue weight and reported as ng/g tissue. Dexamethasone was measured using the same targeted LC–MS/MS workflow adapted for exogenous glucocorticoid detection, as previously described for LC–MS/MS-based dexamethasone quantification^80^.

### Bulk RNA sequencing of primary GBM samples

Total RNA was extracted from tumour tissue using RNeasy Plus Mini Kit, Qiagen according to the manufacturer’s instructions. Libraries were prepared using TruSeq Stranded mRNA Library Prep, Illumina and sequenced on an Illumina NovaSeq 6000 to generate paired-end 150-bp reads. Reads were trimmed and aligned to the human reference genome GRCh38. Then gene-level counts were generated. Downstream analyses were performed in R.Counts were normalised using DESeq2 or converted to transcripts per million (TPM) where appropriate. Expression of steroid receptor genes, including NR3C1, AR, ESR1, ESR2 and PGR, was extracted from the normalised matrix for comparison across samples.

### PBMC isolation and generation of patient-derived monocyte-derived DCs

Fresh peripheral blood mononuclear cells (PBMCs) were isolated from buffy coats or peripheral blood samples from GBM patients by density-gradient centrifugation as previously described^34^. CD14^+^ monocytes were enriched using CD14 MicroBeads (Miltenyi Biotec), yielding >96% CD14+ purity as assessed by flow cytometry. Purified monocytes were differentiated into monocyte-derived dendritic cells (moDCs) for 7 d in RPMI 1640 medium supplemented with 10% heat-inactivated FBS, GM-CSF (50 ng ml−1; PeproTech) and IL-4 (10 ng ml−1; PeproTech)^81^. Where indicated, glucocorticoid-treated DCs were generated by adding dexamethasone (100 nM; Sigma-Aldrich/Merck) 24 h before maturation, whereas control DCs were cultured without dexamethasone. For maturation, LPS (100 ng ml−1) was added 24 h before cell harvest. Mature DCs were collected for downstream flow cytometry, transcriptomic analysis, functional assays or cytokine profiling.

### RNA extraction and sequencing in patient-derived DCs

For dendritic cell RNA-seq experiments, RNA was extracted using the RNeasy Plus Mini Kit (QIAGEN). RNA concentration and quality were assessed before library preparation using fluorometric quantification and electrophoretic fragment analysis, such as Qubit and Bioanalyzer/TapeStation. Libraries were checked by Qubit, real-time PCR and Bioanalyzer-based size-distribution analysis before pooling. Libraries were quantified using Qubit and real-time PCR, and fragment-size distribution was assessed using an Agilent Bioanalyzer. Quantified libraries were pooled at equimolar ratios and sequenced by Novogene on an Illumina NovaSeq 6000 platform to generate 150-bp paired-end reads.

### Bulk RNA-seq analysis

Raw sequencing reads were assessed using FastQC and aligned to the human genome hg38 using HISAT2. SAM files were converted and sorted into BAM files using Samtools. Gene-level counts were generated using htseq-count. Differential expression analysis was performed in R using DESeq2 version 1.38.3 ^82^. Principal component analysis was used to assess sample-level transcriptional structure and treatment-associated separation. Gene set enrichment analysis was performed using GSEA ^83^ and clusterProfiler version 4.6.2 ^84^. Pathways related to T cell functions, glucocorticoid response, antigen presentation, co-stimulation, inflammatory cytokine production, dendritic cell maturation and immune suppression were sourced from the Molecular Signatures Database (MsigDB) (http://www.gsea-msigdb.org/gsea/msigdb/index.jsp). RNA-seq immune deconvolution was performed using the CIBERSORT with the LM22 signature matrix ^85,86^.

### ATAC-seq in patient-derived DCs

Cells were washed twice in ice-cold PBS and lysed in 100 μl ATAC lysis buffer on ice. Following centrifugation cells were resuspended in 50 μl tagmentation mix containing tagmentation buffer, 1% digitonin, Tween-20 with assembled Tn5 transposomes (Active Motif). Tagmentation was performed at 37 °C for 30 min with agitation at 800 rpm. DNA was purified using silica columns (Active Motif) according to the manufacturer’s instructions and eluted in 35 μl elution buffer. Libraries were amplified using Q5 polymerase (New England Biolabs) with indexed Nextera-compatible primers following an initial extension at 72 °C for 5 min and ten cycles of PCR (98 °C for 10 s, 63 °C for 30 s and 72 °C for 1 min). Amplified libraries were double size selected and purified using SPRI beads, eluted in 20 μl elution buffer and quantified using fluorometric assays. Library size distributions were assessed using a TapeStation (Agilent Technologies) before sequencing

### NR3C1 CUT&Tag in patient-derived DCs

CUT&Tag was performed using a modified version of the protocol described by Kaya-Okur et al. ^46^. Patient-derived dendritic cells were harvested, mildly crosslinked with 0.1% formaldehyde in PBS using formaldehyde solution (Thermo Fisher Scientific), quenched with glycine, washed and bound to concanavalin A-coated magnetic beads (Bangs Laboratories). Bead-bound cells were incubated overnight at 4°C with anti-NR3C1/GR antibodies (Cell Signaling Technology, Invitrogen) or rabbit IgG control (Abcam), followed by secondary antibody incubation using guinea pig anti-rabbit IgG (Antibodies Online). Loaded pA-Tn5 transposase (Diagenode) was then added, and tagmentation was activated by MgCl₂ at 37°C. DNA fragments were recovered after reverse crosslinking with EDTA, SDS and Proteinase K (New England Biolabs), followed by phenol–chloroform–isoamyl alcohol extraction (Sigma-Aldrich), chloroform purification and ethanol precipitation using sodium acetate (Invitrogen) and GlycoBlue (Invitrogen). Sequencing libraries were generated using NEBNext High-Fidelity 2× PCR Master Mix (New England Biolabs), with additional amplification cycles determined by SYBR Green qPCR. Libraries were purified and size-selected using AMPure XP beads (Beckman Coulter) and sequenced on an Illumina NovaSeq X platform.

### CUT&Tag data processing and analysis

Paired-end CUT&Tag reads were trimmed using Trim Galore v0.6.10. Trimmed reads were aligned to the human hg38 reference genome using Bowtie2 (v2.5.1). Primary aligned fragments shorter than 1 kb were converted to BEDPE format using bedtools v2.31.1, filtered against the hg38 blacklist and converted to bedGraph coverage files using bedtools genomecov. Fragment coverage was normalised to fragments per million mapped reads and converted to bigWig format using bedGraphToBigWig v4 for genome-browser visualisation.

NR3C1-bound regions were identified from normalised coverage using SEACR v1.3, with matched IgG pulldown libraries used as controls. Stringent peaks were used for downstream analyses. Motif enrichment analysis was performed using HOMER findMotifsGenome.pl v5.1 and the hg38 motif database. Peaks were annotated using ChIPseeker v1.40.0 with TxDb.Hsapiens.UCSC.hg38.knownGene v3.18.0 and org.Hs.eg.db v3.19.1, considering promoter regions defined as ±3 kb from transcription start sites.

For differential NR3C1 occupancy analysis, CUT&Tag signal within ±3 kb of each GENCODE basic transcript transcription start site was quantified using bigWigAverageOverBed v2. Transcript-level signals were aggregated to protein-coding genes by selecting, for each gene, the transcription start site window with the highest median signal across samples. Log2-transformed signals were compared between dexamethasone-treated and control DCs. Representative NR3C1 occupancy tracks, including loci near C5AR1, NR1H3, JAML and SLC38A2, were visualised using bigWig tracks in a genome browser.

### SB28 glioblastoma model

SB28-Ohlfest murine glioma cells were obtained from DSMZ, catalogue no. ACC 880. SB28 cells were cultured in RPMI medium supplemented with 10% fetal bovine serum (FBS; Life Technologies, Invitrogen) and 100 U ml−1 penicillin–streptomycin. For tumour growth experiments, 0.5 × 106 SB28 cells suspended in sterile PBS were injected subcutaneously into the flank of wild-type C57BL/6 mice, *Itgax*^Cre^, *Nr3c1*^fl/fl^ or *Nr3c1*^fl/fl^;*Itgax*^Cre^ mice. Once tumours became visible, tumour dimensions were measured every other day using callipers. Tumour volume was calculated using the ellipsoid formula: tumour volume = 1/2 × longest diameter × shortest diameter².

For dexamethasone treatment experiments, mice received dexamethasone at 10 mg kg−1 body weight by oral gavage once daily for 10 consecutive days, starting 5–7 d after tumour inoculation. Tumour volumes were monitored until the experimental endpoint.

### SB28 tumour tissue processing

Tumours were excised at endpoint and mechanically dissociated before enzymatic digestion in RPMI medium containing 3% FBS, MgCl₂, 1 mg ml−1 collagenase D (Roche), 1 mg ml−1 collagenase A (Roche) and 0.4 mg ml−1 DNase I (Sigma). Digestion was performed at 37 °C for 30–40 min with gentle agitation. Collagenase activity was stopped by adding EDTA to a final concentration of 5 mM. Digested tissues were passed through 70-μm cell strainers (Falcon) to generate single-cell suspensions for downstream flow cytometry, immune-cell sorting or single-cell RNA-seq analysis.

### Non-viral CRISPR–Cas9 editing of NR3C1 in GBM patients derived dendritic cells

Three end-protected synthetic sgRNAs targeting the human NR3C1 locus were designed and obtained from Synthego for evaluation. Two sgRNAs targeted exon 2: C2.1, targeting 5’-CTTTAAGTCTGTTTCCCCCG-3’, and C2.2, targeting 5’-CATCGAACTCTGCACCCCTG-3’. A third sgRNA, C5, targeted exon 5: 5’-AACCTCCAACAGTGACACCA-3’.

Primary human monocyte-derived dendritic cells (moDCs) were harvested and centrifuged at 300g for 5 min at room temperature in 15-ml Falcon tubes, using acceleration setting 9 and deceleration setting 6. Cells were washed once in 1× PBS and resuspended in 20 µl P3 Primary Cell Nucleofector Solution (Lonza) at approximately 4 × 10^5 cells per nucleofection. For each reaction, CRISPR–Cas9 ribonucleoprotein (RNP) complexes were assembled by combining 4 µg TrueCut Cas9 v2 protein (Thermo Fisher Scientific) with 80 pmol of the relevant sgRNA. A Cas9-only condition without sgRNA was included as a negative control. Electroporation was performed in 20-µl Nucleocuvette strips using the Lonza 4D-Nucleofector with pulse code DJ-108. Immediately after electroporation, cells were transferred into 12-well plates containing 1 ml pre-warmed culture medium and cultured for 3 d before downstream analysis.

For genomic DNA extraction, a 10-µl aliquot of cell suspension, corresponding to approximately 4,000 cells, was transferred to a PCR tube and mixed with 40 µl lysis buffer consisting of 50 mM Tris-HCl pH 8.0, 1 mM EDTA, 0.5% Tween-20 and 100 µg ml−1 proteinase K. Samples were incubated at 55 °C for 15 min, followed by proteinase K inactivation at 85 °C for 10 min.

### Sanger sequencing and indel analysis

Target loci were amplified by PCR in 20-µl reactions containing 2 µl cell lysate, 0.3 µM each forward and reverse primer, and Taq PCR Master Mix (APExBIO). PCR amplification was performed on a Bio-Rad T100 Thermal Cycler using the following cycling conditions: initial denaturation at 94 °C for 3 min; 35 cycles of 94 °C for 30 s, 60 °C for 30 s and 72 °C for 45 s; and final extension at 72 °C for 5 min. Primers were designed to generate amplicons compatible with Sanger trace deconvolution.

For C2.1, the primers were:

forward, 5’-TTCTGCGTCTTCACCCTCAC-3’;

reverse, 5’-ACTGGGGCTTGACAAAACCA-3’.

For C2.2, the primers were:

forward, 5’-GGCGGGAGAAGACGATTCAT-3’;

reverse, 5’-AATCCTCACCGTTGGCCAAT-3’.

For C5, the primers were:

forward, 5’-ACTGTGTAGCGCAGACCTTC-3’;

reverse, 5’-TCACCTGACTCTCCCCTTCA-3’.

PCR products were purified using the QIAquick PCR Purification Kit (Qiagen) and eluted in 30 µl Milli-Q water. DNA concentration was measured using a NanoDrop ND-1000 spectrophotometer. Purified amplicons were diluted to 10 ng µl−1 and submitted for Sanger sequencing by Source BioScience. Sequencing was performed using 3.2 µM primer, with the reverse PCR primer used for C2.1 and the corresponding forward primers used for C2.2 and C5. Editing efficiency and indel composition were determined using Synthego Inference of CRISPR Edits (ICE) analysis, version 1.2, by comparison with Cas9-only control traces.

### Cytokine profiling

Cytokine concentrations in GBM patient DC culture supernatants were measured using the MSD Human Proinflammatory Panel 1, 10-plex assay (Meso Scale Discovery; K15049D-2), which quantifies IFN-γ, IL-1β, IL-2, IL-4, IL-6, IL-8, IL-10, IL-12p70, IL-13 and TNF-α. Samples were clarified by centrifugation, stored at −80 °C and thawed once before analysis.

A total of 25 μl sample, standard or quality-control material was added per well and assayed according to the manufacturer’s protocol. Plates were incubated for 2 h, washed, incubated with detection antibody cocktail for a further 2 h, developed with MSD read buffer and acquired on an MSD S600 instrument. Cytokine concentrations were calculated using MSD Workbench software and reported as pg ml−1. Samples were run in duplicate where sufficient material was available.

### Generation of bone marrow-derived dendritic cells

Femurs and tibias were harvested from control Itgax-Cre mice and Nr3c1fl/fl;Itgax-Cre DC-specific knockout mice. Bone marrow was flushed and processed into single-cell suspensions. Cells were differentiated in RPMI 1640 medium supplemented with 10% FBS, and GM-CSF (10 ng ml−1) for 6–7 d. Non-adherent and loosely adherent cells were harvested as immature BMDCs for antigen-loading and co-culture experiments.

For antigen loading and maturation, bone marrow-derived DCs were treated on day 7 with SB28 tumour-cell lysate together with LPS (100 ng ml^−1^) for 16–24 h. Mature DCs were then harvested and washed thoroughly with sterile PBS before adoptive transfer.

### Preparation of tumour lysates and dendritic-cell loading

Patient-derived GBM cell lines were established from freshly resected primary GBM tissues. Tumour specimens were processed immediately under sterile conditions, rinsed in HBSS and minced into approximately 1-mm³ fragments. Tissue fragments were dissociated with Accutase at 37 °C for 45 min, filtered to obtain single-cell suspensions and treated with red blood cell lysis buffer where required. Cells were then cultured on ECM-coated flasks in serum-free Neurobasal-based complete medium supplemented with B27 without vitamin A, N2, GlutaMAX, penicillin–streptomycin, EGF and FGF. Cultures were maintained at 37 °C with 5% CO₂ and passaged at 80–90% confluency. For human tumour-antigen loading, patient-derived GBM cells were harvested, washed in PBS and counted. Tumour lysates were generated from GBM cells by repeated freeze–thaw cycles, followed by centrifugation to remove cellular debris. Human patient-derived DCs were loaded with autologous GBM tumour lysate at a defined tumour-cell-to-DC ratio of 3:1 for 24 h before the maturation, washed thoroughly and then co-cultured with T cells from the same patient for functional assays.

For mouse experiments, SB28 tumour lysate was generated from cultured SB28 glioma cells using the same cell-number-based approach. Briefly, SB28 cells were harvested, washed, counted and subjected to repeated freeze–thaw cycles in PBS. Lysates were clarified by centrifugation, and bone marrow-derived DCs from control or *Nr3c1*^fl/fl^;*Itgax*^Cre^ mice were loaded with SB28 lysate at a defined tumour-cell-to-DC ratio of 3:1 together with LPS maturation 24 h later. Loaded DCs were washed extensively before use in adoptive-transfer vaccination experiments.

### Intraperitoneal adoptive transfer of dendritic cells

For adoptive transfer experiments, 1 × 10^6 mature DCs were resuspended in 100 μl sterile PBS and injected intraperitoneally into the lower right abdominal quadrant of each mouse. DCs were administered on days 7 and 14 after SB28 tumour implantation. Tumour growth was monitored as described above until the experimental endpoint.

### OT-I antigen-presentation assay

Control or Nr3c1-deficient BMDCs were pulsed with SIINFEKL peptide, 1 μg ml^−1^ for 1–2 h, washed and co-cultured with purified CD8^+^ T cells isolated from spleens of OT-I mice. OT-I CD8^+^ T cells were labelled with CellTrace Violet (CTV; Thermo Fisher Scientific) according to the manufacturer’s instructions before co-culture. After 72 h, T-cell proliferation was assessed by CTV dilution and activation was quantified by expression of CD69 and CD44 using flow cytometry.

### Spectral flow cytometry

For intracellular cytokine staining, single-cell suspensions were stimulated for 4 h with PMA (50 ng ml⁻¹) and ionomycin (500 ng ml⁻¹), with monensin (BioLegend) added during the final 3 h. Cells were stained using standard surface and intracellular staining protocols. Briefly, cells were first stained with LIVE/DEAD Fixable Dead Cell Stain (Thermo Fisher Scientific), followed by surface antibody staining with Bright Staining buffer. Cells were then fixed using eBioscience IC Fixation Buffer and permeabilised with 1× permeabilisation buffer for intracellular cytokine. Fluorochrome-conjugated antibodies were applied as indicated, and cells were washed in PBS containing 3% FCS before acquisition.

Samples were acquired on a Cytek Aurora 5-laser spectral flow cytometer, and data were analysed using FlowJo v10.2. Antibodies used for human DC analyses included CD14–FITC (M5E2, BioLegend, 301804; 1:100), CD86–PE (IT2.2, BioLegend, 305406; 1:200), IFNγ–Alexa Fluor 700 (4S.B3, BioLegend, 502520; 1:100) and TNFα–PE–Cy7 (MAb11, BioLegend, 502930; 1:100). For mouse tumour and DC-vaccine experiments, antibodies included CD45–BV711 (30-F11, BioLegend, 103147; 1:1,000), CD45–BUV563 (30-F11, BD Biosciences, 612924; 1:1,000) or CD45–FITC (30-F11, BioLegend, 103108; 1:1,000); TCRβ–FITC (H57-597, BioLegend, 109206; 1:400) or TCRβ–eFluor 450 (H57-597, Thermo Fisher Scientific/eBioscience, 48-5961-82; 1:400); CD11b–APC–Cy7 (M1/70, BioLegend, 101226; 1:400); CD4–APC (GK1.5, BioLegend, 100412; 1:400) or CD4–PE–Cy7 (GK1.5, BioLegend, 100422; 1:400); CD8α–APC–eFluor 780 (53-6.7, Thermo Fisher Scientific/eBioscience, 47-0081-82; 1:400); CD44–APC (IM7, BioLegend, 103012; 1:400); CD69–PE–Cy5 (H1.2F3, BioLegend, 104510; 1:400); MRC1/CD206–APC (C068C2, BioLegend, 141708; 1:400); IFNγ–PerCP–Cy5.5 (XMG1.2, BioLegend, 505822; 1:200); and TNFα–PE–eFluor 610 (MP6-XT22, Thermo Fisher Scientific/eBioscience, 61-7321-82; 1:200). Fc receptors were blocked using human Fc-blocking solution (BioLegend, 422301) or anti-mouse CD16/32 (clone 93, BioLegend, 101320), according to the manufacturer’s instructions. Dead cells were excluded using LIVE/DEAD Fixable Violet staining.

### Immune deconvolution and correlation analyses

To estimate immune population abundance from bulk GBM RNA-seq data, deconvolution was performed using CIBERSORT^87^. Activated DC abundance and other immune scores were correlated with steroid receptor expression using Pearson’s rank correlation unless otherwise stated. P values were adjusted for multiple comparisons using the Benjamini–Hochberg method where several receptor and cell type pairs were tested simultaneously.

### Single-cell RNA-seq library preparation and sequencing

Tumours were minced and digested as mentioned above. Cell suspensions were filtered through a 70-μm strainer, washed and subjected to red blood cell lysis if necessary. CD45^+^ immune cells were enriched by FACS before counting and single-cell library preparation. Single-cell RNA-seq libraries were prepared at the Cancer Research UK Cambridge Institute Genomics Core Facility using the Chromium Next GEM Single Cell 5’ Kit v2 and associated reagents and protocols from 10x Genomics. Sorted cells were collected in PBS containing 0.04% BSA and loaded onto Chromium microfluidic chips for single-cell partitioning. Gel-bead emulsions were generated on the Chromium Controller using 10x Genomics 5’ v2 chemistry, targeting approximately 10,000 cells per sample.

Barcoded RNA was reverse-transcribed in emulsion using a C1000 Touch Thermal Cycler (Bio-Rad). Resulting cDNA was recovered and assessed for quality and quantity using an Agilent TapeStation 4200. Approximately 1,000 ng cDNA was used for library preparation, with sample-indexing PCR cycles adjusted according to cDNA yield. Final libraries were assessed using the Agilent TapeStation 4200 and quantified using a BMG LABTECH Clariostar Monochromator Microplate Reader for double-stranded DNA measurement. Libraries were normalised to 10 nM, pooled and sequenced on an Illumina NovaSeq 6000, with each pool occupying approximately 16% of a sequencing lane. Sequencing was performed using 10x Genomics-compatible read configurations for 5’ libraries, with the aim of achieving high sequencing depth per cell and approximately 2 billion total reads across the experiment. Base-call files were converted to FASTQ files, and sequencing quality was summarised using MultiQC.

### Single-cell data processing

Raw single-cell RNA-seq data were processed using Cell Ranger version 8.0.1. Reads were demultiplexed, aligned to the appropriate reference genome and summarised into gene–cell UMI count matrices. Genes expressed in fewer than three cells were removed before downstream analysis. For mouse scRNA-seq analysis, cells were retained if they expressed more than 200 and fewer than 8,000 genes, contained fewer than 100,000 UMIs, and had less than 15% mitochondrial gene content. Data were normalised using the LogNormalize method in Seurat version 5.0.3^88^ with a scale factor of 10,000. Mitochondrial gene percentage and total UMI count were regressed out during scaling. Principal component analysis was performed using the top 2,000 highly variable genes identified using Seurat. Shared nearest-neighbour graphs were constructed using FindNeighbors, and unsupervised clustering was performed using FindClusters with resolution 0.6. UMAP and t-SNE visualisations were generated using RunUMAP and RunTSNE. Public and our previously published GBM patient sc/snRNA-seq datasets were obtained from GSE182109^38^ and GBM-Space (https://www.gbmspace.org/)^89^. Gene-set scores were calculated using AddModuleScore in Seurat version 5.0.3. Curated gene sets included M1 and M2 macrophage signatures, GO biological process antigen processing and presentation, Hallmark inflammatory response, GO biological process dendritic cell differentiation, and GO biological process regulation of lymphocyte cytotoxicity, all obtained from the MsigDB (http://www.gsea-msigdb.org/gsea/msigdb/index.jsp).

## Statistical analysis

Statistical analyses were performed using GraphPad Prism [v10] and R [v4.X]. All tests were two-sided unless otherwise stated. For comparisons between two groups, unpaired two-tailed Student’s t-tests were used for normally distributed data and Mann–Whitney U-tests for non-parametric data. Paired analyses used paired t-tests or Wilcoxon signed-rank tests as appropriate. For comparisons involving more than two groups, one-way or two-way ANOVA with post hoc multiple-comparison correction was used. Tumour growth curves were analysed using mixed-effects models or two-way repeated-measures ANOVA, as appropriate. Correlation analyses used Pearson’s rank correlation unless otherwise specified. Differential expression and chromatin analyses were corrected for multiple testing using the Benjamini–Hochberg method. Adjusted P < 0.05 was considered statistically significant. Data are presented as mean ± s.e.m. unless otherwise indicated. Centre lines, boxes and whiskers in box plots represent [median, interquartile range and 1.5 × IQR] or should be defined in the relevant figure legends.

## Acknowledgements

We would like to thank Prof Louise Boyle for sharing Cd11c-Cre mouse line (*Itgax*^Cre^); Joana Cerveira and the entire flow cytometry team at the Department of Pathology for help with flow cytometry; UBS animal facility, Gurdon Institute, for their technical help and animal husbandr; Scientific core at the CRUK Cambridge Institute for HPC resources, and the Genomics core for sequencing services; Silvia Corriero for helful advice on CUT&Tag experiments.We dedicate this work to the memory of our late colleague Professor Gregory J. Hannon.

## Author contributions

Q.Z. and J.P. contributed equally to this work. R.M. and B.M. jointly supervised this study.

Conceptualization: B.M., Q.Z., R.M. and J.P.; Methodology: Q.Z. and J.P.; Investigation: Q.Z., J.P., A.J., E.F., S.C., Y.G., S.G., M.P.P.-C., M.S., A.A.-d., M.P., H.H., C.J.W., G.G., N.Z.M.H. and B.C.N.; Formal analysis: Q.Z., and J.P., with contributions from A.J., E.F., S.C., Y.G., S.G., M.P.P.-C., M.S., A.A.-d., M.P., H.H., C.J.W., G.G., N.Z.M.H. and B.C.N.; Data curation: Q.Z.; Writing – original draft: Q.Z. with input and support from co-authors; Writing – review & editing: B.M. R.M. and other co-authors; Funding acquisition: B.M. and R.M.; Project administration: Q.Z. and J.P.; Supervision: R.R. supervised A.A.-d. and B.M. and R.M managed the team and supervised the study.

## Data and materials availability

The RNA-seq and CUT&Tag-seq datasets generated in this study will be deposited in the Gene Expression Omnibus (GEO) during peer review and will be made publicly available upon acceptance of the peer-reviewed article. Accession numbers will be added to the manuscript once available.

