## Supplementary Figures S1-S4 for "Targeting glucocorticoid signalling in dendritic cells for glioblastoma treatment"

### Supplementary Figure Legends

**Figure S1. Steroid-receptor and glucocorticoid-response gene expression in GBM immune cells**

(A) t-SNE projection of immune cells from a public human glioma scRNA-seq dataset, showing the major myeloid, T-cell and B-cell compartments. (B) Feature plots showing the expression of the glucocorticoid receptor NR3C1, mineralocorticoid receptor NR3C2, oestrogen receptors ESR1 and ESR2, progesterone receptor PGR and androgen receptor AR across the immune-cell embedding. Colour intensity indicates normalised expression. (C) Dot plot showing expression of the canonical glucocorticoid-responsive gene TSC22D3 across annotated GBM myeloid-cell populations. Dot size represents the percentage of cells expressing TSC22D3, and colour indicates average scaled expression. AP, antigen-presentation; DC, dendritic cell; MDSC, myeloid-derived suppressor cell; scRNA-seq, single-cell RNA sequencing; t-SNE, t-distributed stochastic neighbour embedding.

**Figure S2. Integrated epigenomic and transcriptional responses to dexamethasone in patient-derived DCs**

(A) Integrated comparison of dexamethasone-induced changes in TSS-centred NR3C1 CUT&Tag signal and RNA expression. Genes with concordant increases are shown in red, genes with concordant decreases in blue, genes changing in opposite directions in brown, and other overlapping genes in grey; representative genes are labelled. (B) Principal component analysis of ATAC-seq profiles from control and dexamethasone-treated GBM patient-derived DCs. Each point represents one sample, and colours indicate treatment group. (C) Volcano plot of differential chromatin accessibility in dexamethasone-treated versus control DCs. Dexamethasone-open regions are shown in red, dexamethasone-closed regions in blue and non-significant regions in grey; representative peak-associated genes are labelled. Dashed lines indicate the fold-change and adjusted-P-value thresholds used for differential analysis. ATAC-seq, assay for transposase-accessible chromatin using sequencing; Ctrl, control; DC, dendritic cell; Dex, dexamethasone; TSS, transcription start site.

**Figure S3. Effects of DC-specific Nr3c1 deletion on SB28 tumour growth and tumour-infiltrating immune states**

(A) SB28 tumour growth in control and DC-specific Nr3c1 conditional-knockout (Nr3c1cKO) mice in the absence of dexamethasone ($n = 7$ mice per group). Data are presented as mean +/- s.e.m. (B) Endpoint tumour volume for the experiment shown in A. Each point represents one mouse; bars show mean +/- s.e.m. The modest reduction in tumour volume did not reach statistical significance ($P = 0.0721$). (C) t-SNE projection of re-clustered tumour-infiltrating T cells from dexamethasone-treated control and Nr3c1cKO tumours, resolving naive T cells, central-memory T cells, cytotoxic CD8+ T cells, conventional CD4+ T cells, regulatory T cells, gamma-delta T17 cells and IFN-high T cells. (D) Dot plot of canonical marker-gene expression used to annotate the T-cell states shown in C. (E) Gene set enrichment analysis of tumour-infiltrating CD8+ T cells from dexamethasone-treated Nr3c1cKO versus control tumours, showing positive enrichment of HALLMARK_INTERFERON_ALPHA_RESPONSE (nominal $P = 0.001595$; NES = 1.93) and HALLMARK_INTERFERON_GAMMA_RESPONSE (nominal $P = 0.001473$; NES = 1.58) in the Nr3c1cKO group. (F) Dot plot of canonical marker-gene expression used to annotate 15 re-clustered myeloid states, including Abca1+, Mertk+, Lpl+, Folr2+ and Arg1+ tumour-associated macrophages, antigen-presentation-high macrophages, inflammatory and IFN-high monocytes, S100A8+ monocytic MDSCs, pDCs and cDC subsets. In D and F, dot size represents the percentage of cells expressing each gene and colour indicates average scaled expression. AP, antigen-presentation; cDC, conventional dendritic cell; IFN, interferon; MDSC, myeloid-derived suppressor cell; NES, normalised enrichment score; pDC, plasmacytoid dendritic cell; TAM, tumour-associated macrophage; Tcm, central-memory T cell; Treg, regulatory T cell.

**Figure S4. *NR3C1* deletion reverses dexamethasone-associated regulatory programmes in patient-derived DCs**

(A) Four-way overlap among genes linked to dexamethasone-regulated ATAC-seq regions, dexamethasone-regulated NR3C1 CUT&Tag regions, dexamethasone-responsive RNA-seq genes and genes differentially expressed following NR3C1 deletion under dexamethasone exposure. A total of 198 genes were shared across all four datasets. (B) Heat map showing log2 fold changes for 48 representative genes from the 198-gene intersection across the four datasets. Established glucocorticoid-responsive or restrained-state genes, including FKBP5, TSC22D3, IL1RL1, PIK3IP1, C5AR1 and MAFB, were induced by dexamethasone and reduced after NR3C1 deletion, whereas DC activation-associated genes, including CD80, CD86, IL12A, IL12B, CCR7, CD83, BATF3 and JAML, showed the reciprocal pattern. (C) Gene set enrichment analysis of NR3C1-edited versus Cas9-control DCs under dexamethasone exposure, showing enrichment of antigen processing and presentation of endogenous antigen (nominal P = 0.0132; NES = 1.68) and inflammatory response to antigenic stimulus (nominal P = 0.0122; NES = 1.55). ATAC-seq, assay for transposase-accessible chromatin using sequencing; DC, dendritic cell; NES, normalised enrichment score.

Figure S1

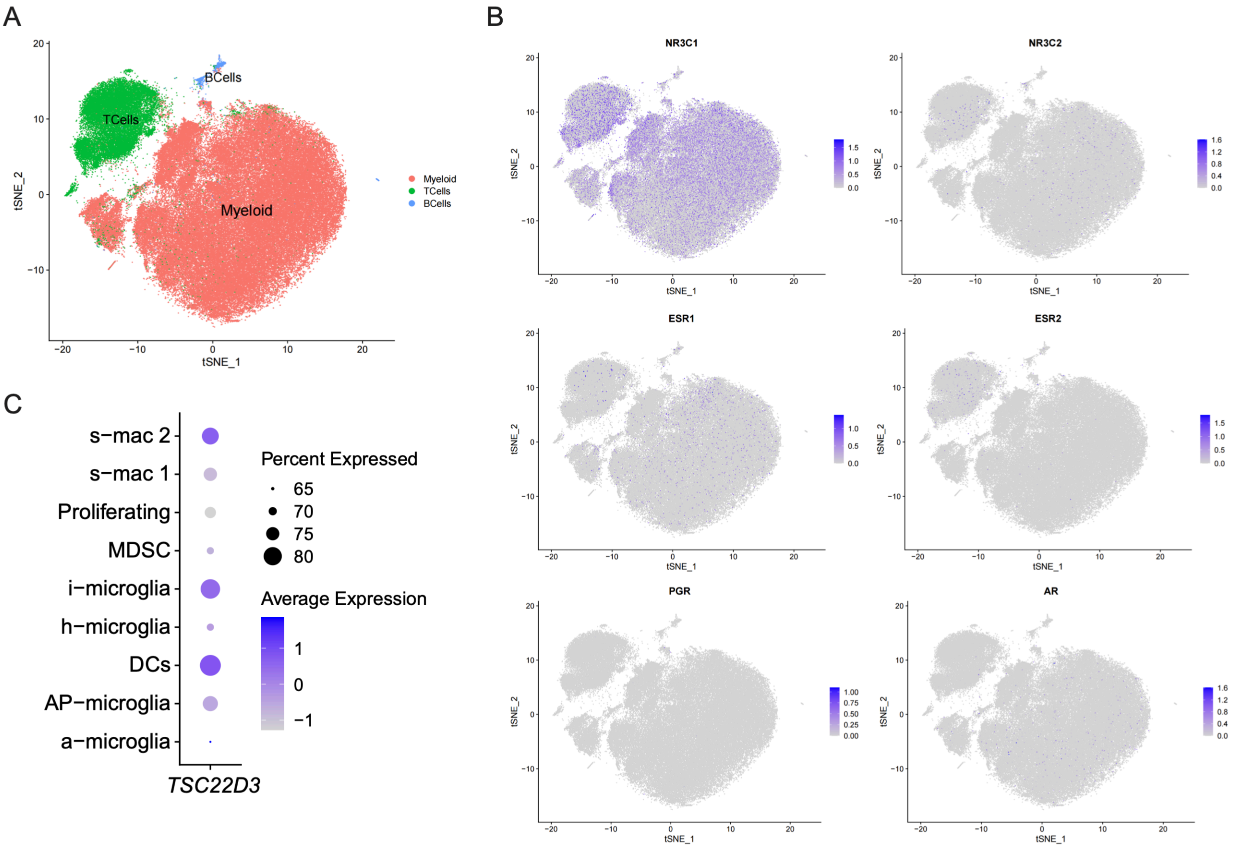

A
B
C
BCells
TCells
Myeloid
tSNE_1
tSNE_2
Myeloid
TCells
BCells
NR3C1
NR3C2
ESR1
ESR2
PGR
AR
s−mac 2
s−mac 1
Proliferating
MDSC
i−microglia
h−microglia
DCs
AP−microglia
a−microglia
TSC22D3
Percent Expressed
65
70
75
80
Average Expression
1
0
−1

Figure S2

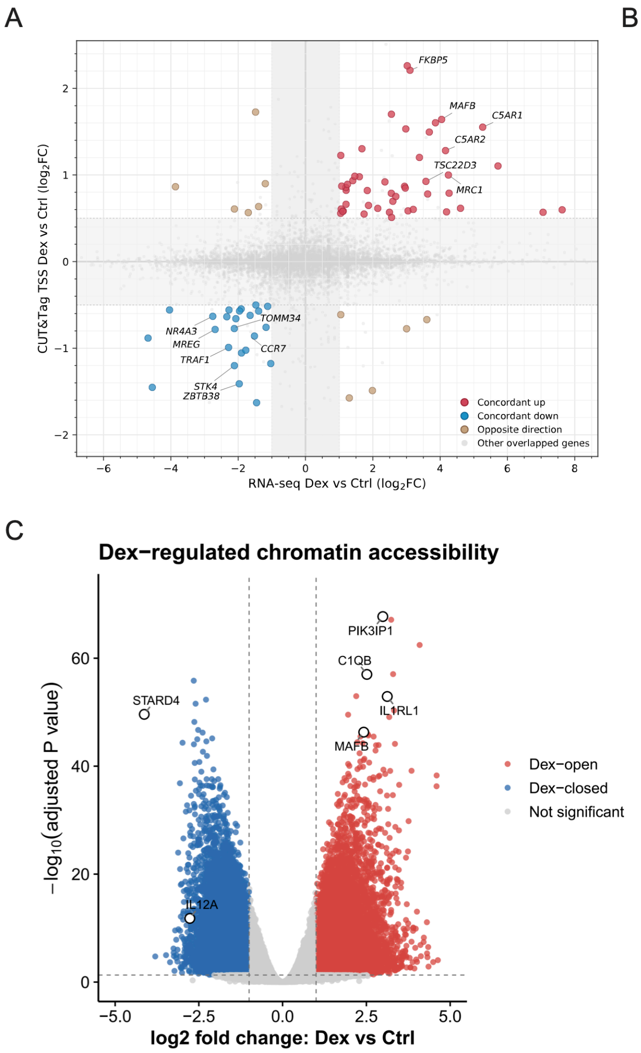
A
FKBP5
MAFB
C5AR1
C5AR2
TSC22D3
MRC1
TOMM34
NR4A3
MREG
CCR7
TRAF1
STK4
ZBTB38
Concordant up
Concordant down
Opposite direction
Other overlapped genes
CUT&Tag TSS Dex vs Ctrl (log2FC)
RNA-seq Dex vs Ctrl (log2FC)
B
PC2: 31% variance
PC1: 45% variance
Ctrl
Dex
C
Dex−regulated chromatin accessibility
PIK3IP1
C1QB
STARD4
IL1RL1
MAFB
IL12A
−log10(adjusted P value)
log2 fold change: Dex vs Ctrl
Dex−open
Dex−closed
Not significant

Figure S3

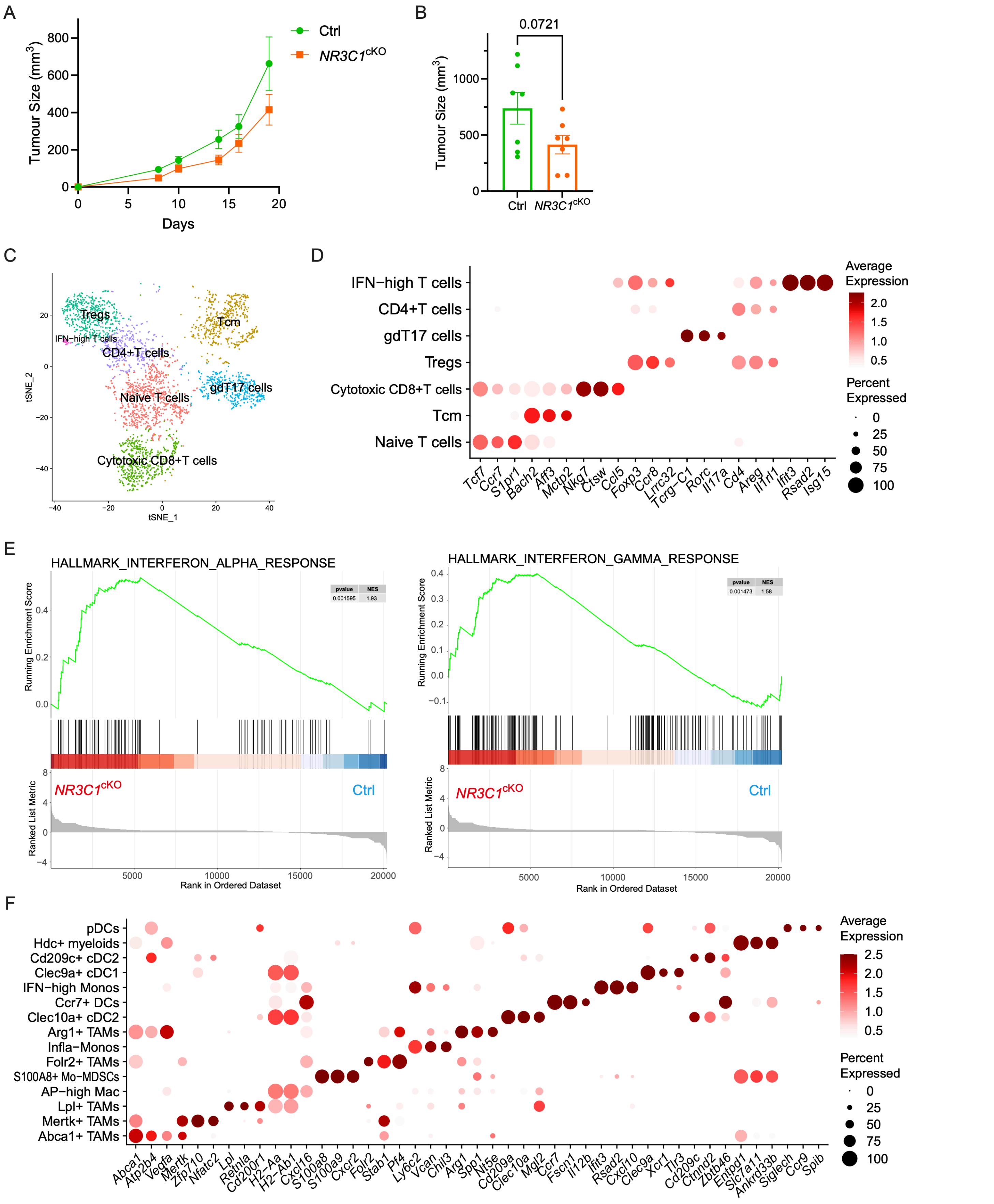
A
Ctrl
NR3C1cKO
Tumour Size (mm3)
800
600
400
200
0
0
5
10
15
20
Days
B
0.0721
Tumour Size (mm3)
1000
500
0
Ctrl
NR3C1cKO
C
Tregs
IFN−high T cells
CD4+T cells
Tcm
gdT17 cells
Naive T cells
Cytotoxic CD8+T cells
tSNE_2
tSNE_1
D
IFN−high T cells
CD4+T cells
gdT17 cells
Tregs
Cytotoxic CD8+T cells
Tcm
Naive T cells
Tcf7
Ccr7
S1pr1
Bach2
Aff3
Mctp2
Nkg7
Ctsw
Ccl5
Foxp3
Ccr8
Lrrc32
Tcrg−C1
Rorc
Il17a
Cd4
Areg
Il1rl1
Ifit3
Rsad2
Isg15
Average Expression
2.0
1.5
1.0
0.5
Percent Expressed
0
25
50
75
100
E
HALLMARK_INTERFERON_ALPHA_RESPONSE
pvalue
NES
0.001595
1.93
Running Enrichment Score
Ranked List Metric
NR3C1cKO
Ctrl
Rank in Ordered Dataset
HALLMARK_INTERFERON_GAMMA_RESPONSE
pvalue
NES
0.001473
1.58
Running Enrichment Score
Ranked List Metric
NR3C1cKO
Ctrl
Rank in Ordered Dataset
F
pDCs
Hdc+ myeloids
Cd209c+ cDC2
Clec9a+ cDC1
IFN−high Monos
Ccr7+ DCs
Clec10a+ cDC2
Arg1+ TAMs
Infla−Monos
Folr2+ TAMs
S100A8+ Mo−MDSCs
AP−high Mac
Lpl+ TAMs
Mertk+ TAMs
Abca1+ TAMs
Abca1
Atp2b4
Vegfa
Mertk
Zfp710
Nfatc2
Lpl
Retnla
Cd200r1
H2−Aa
H2−Ab1
Cxcl16
S100a8
S100a9
Cxcr2
Folr2
Stab1
Pf4
Ly6c2
Vcan
Chil3
Arg1
Spp1
Nt5e
Cd209a
Clec10a
Mgl2
Ccr7
Fscn1
Il12b
Ifit3
Rsad2
Cxcl10
Clec9a
Xcr1
Tlr3
Cd209c
Ctnnd2
Zbtb46
Entpd1
Slc7a11
Ankrd33b
Siglech
Ccr9
Spib
Average Expression
2.5
2.0
1.5
1.0
0.5
Percent Expressed
0
25
50
75
100

Figure S4

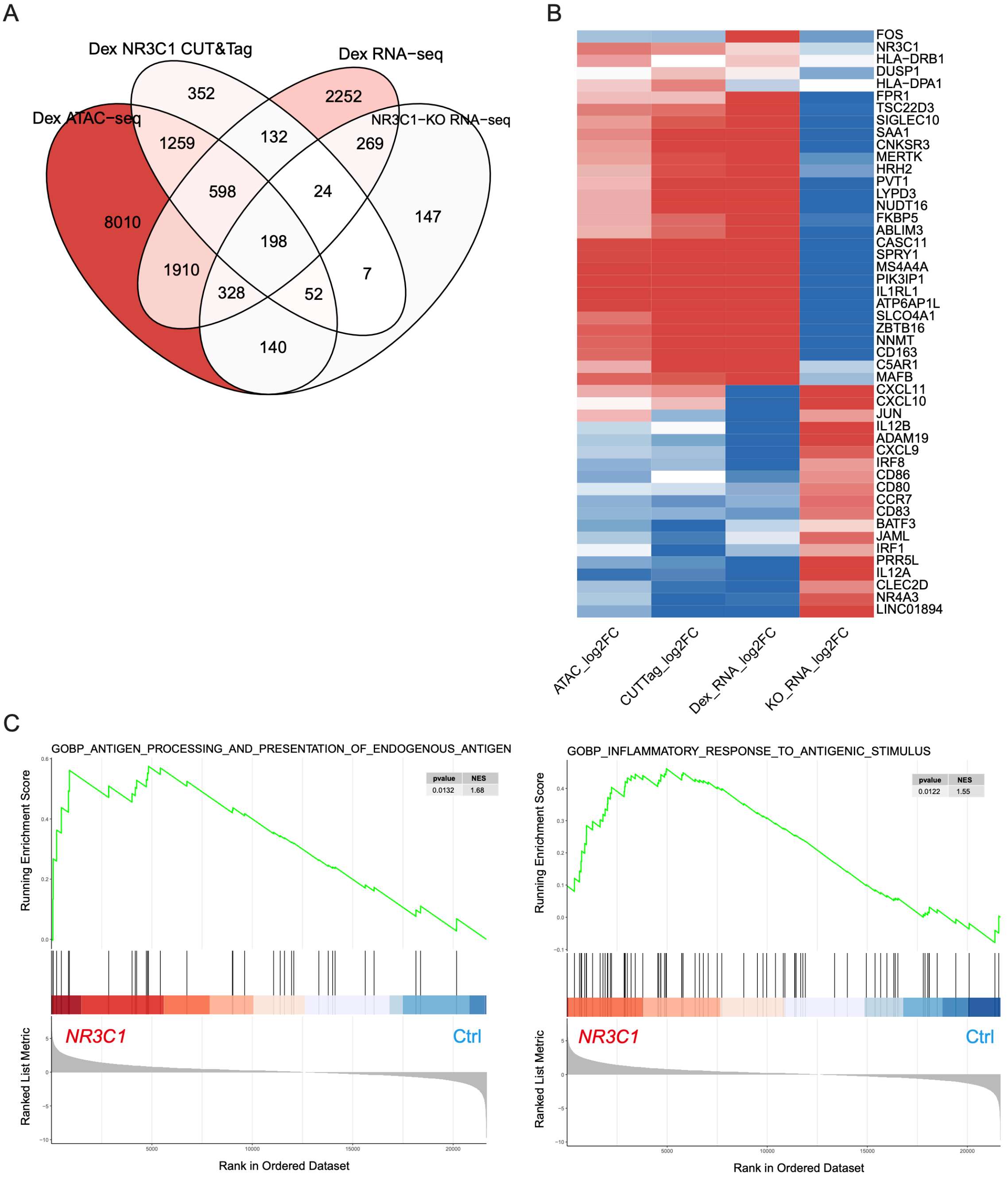
A
Dex NR3C1 CUT&Tag
Dex RNA−seq
Dex ATAC−seq
NR3C1−KO RNA−seq
352
2252
1259
132
269
598
24
8010
198
147
1910
328
52
7
140
B
FOS
NR3C1
HLA−DRB1
DUSP1
HLA−DPA1
FPR1
TSC22D3
SIGLEC10
SAA1
CNKSR3
MERTK
HRH2
PVT1
LYPD3
NUDT16
FKBP5
ABLIM3
CASC11
SPRY1
MS4A4A
PIK3IP1
IL1RL1
ATP6AP1L
SLCO4A1
ZBTB16
NNMT
CD163
C5AR1
MAFB
CXCL11
CXCL10
JUN
IL12B
ADAM19
CXCL9
IRF8
CD86
CD80
CCR7
CD83
BATF3
JAML
IRF1
PRR5L
IL12A
CLEC2D
NR4A3
LINC01894
ATAC_log2FC
CUTTag_log2FC
Dex_RNA_log2FC
KO_RNA_log2FC
C
GOBP_ANTIGEN_PROCESSING_AND_PRESENTATION_OF_ENDOGENOUS_ANTIGEN
Running Enrichment Score
pvalue NES
0.0132 1.68
Ranked List Metric
NR3C1
Ctrl
Rank in Ordered Dataset
GOBP_INFLAMMATORY_RESPONSE_TO_ANTIGENIC_STIMULUS
Running Enrichment Score
pvalue NES
0.0122 1.55
Ranked List Metric
NR3C1
Ctrl
Rank in Ordered Dataset
